# RELAX does not reproduce its own estimates at default settings, and its output does not show it

**DOI:** 10.64898/2026.09.20.751138

**Authors:** MinSeo Kim, Jae-Ho Shin

## Abstract

Selection-intensity estimates from RELAX are reported as a point value of *K* with a likelihood-ratio *P*. We report that, at default settings and on data of ordinary size, the program does not reproduce its own fits. Of 27 enzyme entries refitted under two optimiser configurations, none reproduced its log-likelihood to within 0.01 units; the median change was 103 units, the largest over 3,400, and four verdicts reversed. Eighty null orthologues reproduced none. A byte-identical command returned a distinct likelihood on every repetition, single-threaded, across three releases, and on alignments simulated under the fitted model, where 3.3 per cent of replicates reproduced. The documented random-number seed never reaches the generator when assigned on the command line, yet reads back as the value supplied. PAML localises the cause: its two-ratio model, without site classes, reproduced its log-likelihood for all 288 genes; its site-class models agreed for 27 to 67 per cent. The instability follows the mixture over sites, not the program. The output does not show it: 46 of 410 fits ended with a negative likelihood-ratio statistic, impossible under convergence, and 123 of 410 report a *K* re-estimated under a domain restriction rather than the unconstrained maximum. Of 234 published studies using RELAX, none reported a seed. Seeding while holding the thread count at one reproduced sixty of sixty runs on twenty genes under two releases; the seed alone reproduced none of five, and no documentation states the second condition. We recommend that fits be repeated and their dispersion published.

## INTRODUCTION

RELAX (Wertheim *et al*., 2015) tests whether selection on a set of branches is relaxed or intensified relative to the rest of a phylogeny, and it has become the standard implementation of that comparison. The analyst partitions the tree into a test set and a reference set; the program fits a three-class distribution of ω to the reference branches and a single exponent, *K*, that raises or flattens that distribution on the test branches; and the result is reported as an estimate of *K* with a likelihood-ratio *P*. Because the method needs no experimental data and converts a categorical trait directly into a hypothesis about selection, it is applied widely, usually once per gene, at the program’s default settings.

An estimate of this kind rests on an optimisation, and what an analyst can learn about that optimisation from its output is limited. Convergence is not reported as a property the user can verify, the likelihood surface is not shown, and the parameter that is returned looks the same whether the search that produced it ended at a maximum or somewhere else. Reproducibility is the one check that requires nothing beyond the program itself: the same command on the same data should return the same fit. We ran that check, and this manuscript reports what it returned.

The occasion was our own analysis of plant-cell-wall-degrading enzyme genes across ectomycorrhizal fungi, in which relaxed selection on several families had been reported on the basis of RELAX fits (Kim and Shin, 2026). A companion manuscript (Kim and Shin, in preparation) shows that the type I error rate of that design is set by where the labels are placed, and withdraws the claims on that ground. The present paper documents a second failure, independent of the first and found before it: the fits did not reproduce. None of 27 enzyme entries returned the same log-likelihood when the number of optimiser starting points was changed, none of 80 null orthologues did either, and repeating a byte-identical command gave a different answer every time.

We then asked where the instability comes from, since the answer determines what can be done about it. We excluded the data and the model by simulating alignments under the fitted model and finding the same behaviour on them. We excluded parallel arithmetic by single-threaded execution, which widened the spread. We found that the documented random-number seed has no effect, and why. And we localised the cause to the mixture over sites by fitting the same alignments in a second program, PAML, whose model without site classes reproduced exactly and whose models with site classes did not.

The failure is invisible in ordinary output, and that is the observation we would put first. A run that terminates in a state convergence makes impossible returns a *P* value formatted like any other; a parameter returned exactly at its null value looks like a converged estimate of no effect; and a reported *K* may be a re-estimate under a domain restriction rather than the maximum the output table implies. We quantify each of these from the program’s own diagnostics, ask how the published literature reports RELAX analyses, describe the one route to a reproducible fit that we found, which the documentation does not lead to, and state what we would now require of any such analysis before accepting its result. We accepted our own results for several months, and the checks that eventually overturned them were inexpensive.

## MATERIALS AND METHODS

### M1. Alignments and entries

Every fit reported here was made on material described in full in the companion study, and only what is needed to read the results is repeated. Coding sequences were available for 51 of the 183 genomes of that study’s panel, 21 of them from ectomycorrhizal (ECM) species; codon alignments and gene trees were built as described there (M4). Twenty-eight plant-cell-wall-degrading enzyme entries were tested, twenty families, seven *GH5* subfamilies and one subsampled *GH5* alignment; twenty-seven returned a fit under both optimiser settings compared below, and every count of analysed entries in this paper is that 27. The null sets, built from single-copy orthologues identified with BUSCO 6.1.0 (Manni *et al*., 2021), and the simulated replicates are defined in the following section; all alignments, trees and fitted outputs are deposited (Data availability).

### M2. Tests of selection, their reproducibility, and the null distributions

Selection regimes were tested with RELAX (Wertheim *et al*., 2015) in HyPhy 2.5.100, MP build (Kosakovsky Pond *et al*., 2020); RELAX.bf carries its own version string, 4.7. Terminal branches leading to ECM species were labelled {Test}, leaving non-ECM tips and all internal branches as the reference set. *K* and its likelihood-ratio *P* came from the RELAX JSON, *K* < 1 denoting relaxed and *K* > 1 intensified selection on the test branches; rejections are counted two-sided. Runs used --starting-points 5 unless stated otherwise, with --srv, --multiple-hits and --error-sink off at their defaults, the usage characterised here. Command lines and seeds are in Supplementary Methods S1.

### Reproducibility and convergence diagnostics

Reproducibility was measured first, with identical command lines re-executed sequentially on test genes, comparing *K*, log-likelihood and *P* across replicates, with seed and thread count varied as controls, and then, in a pre-registered extension whose gene list, arms, criterion and predictions were fixed before execution, on twenty null orthologues under four combinations of release, seed and thread count (Supplementary S2.12). The sign of the likelihood-ratio statistic was read from every JSON, and convergence flags were censused over the observed and null-distribution batches only, the 410 fits of Results 2, with the simulation, misspecification-control, ladder, CAZyme and replicate batches analysed separately and excluded from that pool (S1).

We use *arm* throughout for one complete configuration of program, model, flag settings, labelling scheme and starting-point budget, run over a whole gene set. Arms are compared only against their own baseline, and cross-arm comparisons are made on the genes interpretable in every arm compared, with that common set stated. A fit is *interpretable* when it returns a fitted *K* and a non-negative likelihood-ratio statistic; a fit terminating with a negative statistic after its five rescue passes is assigned *P* = 1, cannot reject, and is excluded from any rate quoted on interpretable fits. Zero-byte outputs count as failures, and every paired comparison uses the intersection of genes succeeding in both arms, with its size reported.

### Null distributions

Three null sets were built from the same 51 genomes with coding sequences, so null and test share the alignment pipeline. Two are described below; the third, an earlier random labelling drawn from the whole panel rather than from the non-ectomycorrhizal species and fitted from one starting point rather than five, is null A of the companion study (Results 3) and is reported there only as a check.

#### Null B, ECM labelling on single-copy orthologues

Every single-copy fungi_odb10 BUSCO gene (Manni *et al*., 2021) in all 51 CDS genomes went through the alignment pipeline of the companion study (M4) with the same 21 ECM tips labelled {Test}. Eighty-nine met the presence criterion, one (186574at4751) failing tree inference on a non-rectangular alignment; of 88 submitted, 82 returned an interpretable block. No gene was dropped on the basis of its result.

#### Null C, ECM-free random labelling

The same alignments and gene trees were re-analysed with only labels changed, 21 tips per gene being drawn at random from the 30 non-ECM genomes, the deposited draws rather than the seed defining the analysis (S1). All 88 draws contained zero ECM tips and exactly 21 test tips; 81 returned an interpretable block, a different final *n* on the same alignments. A retired earlier version drew from all 51 genomes (S1).

Tests of a proportion against a fixed value are exact binomial; interval estimates of a proportion are Wilson score intervals (S1).

#### A second implementation

To separate the test from HyPhy’s implementation, the question was repeated in codeml (PAML 4.10.10; Yang, 2007), fitting one-ratio against two-ratio with the labelled branches as foreground, likelihood-ratio test on one degree of freedom, each model fitted from *ω* = 0.5 and 1.5 with the higher likelihood retained. Four sets mirror the RELAX runs. These are null C (88 genes), the *K* = 1 simulated replicates of the shallow set labelled on tips (100) and the same alignments labelled on internal branches (100), and the divergence-matched *K* = 1 replicates labelled on tips (80). The matched set was fitted under terminal labelling only and so has no internal-branch counterpart in this program (S1).

#### A second model family within that program

Because the two-ratio model assumes one *ω* across sites, those sets were analysed again under Clade Model C against its null M2a_rel (Weadick and Chang, 2012), which it exceeds by exactly one parameter. An initial pass truncated the slowest fits at rates differing between observed and simulated sets, so all were re-run to completion; of the divergence-matched set’s 320 clade-model fits, two remained incomplete (S1).

### Simulation under a known generating value

Alignments were simulated under the fitted model and re-analysed with the identical invocation. Each template gene supplied its own parameters, taken from its own RELAX null fit and its own alignment and {Test}-labelled tree, and sequences were evolved with pyvolve 1.1.0 (Spielman and Wilke, 2015) under MG94 (Muse and Gaut, 1994), the reference partition at the fitted *ω* classes and the test partition at *ω*^*K*. Five template genes were simulated at *K* ∈ {0.5, 0.7, 1.0}. No random seed was passed to the simulator for this first set, so its replicates cannot be regenerated and are deposited individually (S1).

A second, divergence-matched set was built because the first was far shallower than the alignments it stood for. Branch-length scale was set per template by log-scale bisection until the simulated alignment matched that template’s observed mean pairwise identity, separately at each generating *K*; one template could not reach its target and was dropped. Only the scale is altered, so the generating *K* is preserved.

Every simulated alignment was fitted at --starting-points 1 and --starting-points 5 into separate output sets, the five-point arm being the one compared against the observed nulls. A third arm labelled internal branches rather than tips, re-using those alignments, which at *K* = 1 are independent of the labelling (S1).

#### Reporting practice in the published literature

To ask whether the failures this paper reports are visible to a reader of the literature, Europe PMC was searched over full text for records using RELAX, and the retrieved full texts were scanned for a stated random-number seed, a stated number of optimiser starting points, and repeated fitting undertaken to check reproducibility. Because RELAX is also an ordinary English word, a record was counted only where the token co-occurred with unambiguous evidence of the program. The accounting from search to denominator, the validation of the search, and the script itself are given in Supplementary S2.13.

## RESULTS 1 — THE FITTED VALUES ARE NOT REPRODUCIBLE, AND THE CAUSE IS THE OPTIMISER

### The comparison is not confounded

Eighty null genes were fitted twice under settings that differed only in the number of optimiser starting points (one versus five). Before interpreting any difference we verified that nothing else did: alignment file, sequence and codon-site counts, tree string and {Test}/{Reference} partition were identical in all eighty pairs, and separately in the twenty-seven paired enzyme entries; the estimated parameter count agreed in seventy-nine of eighty; and every fit came from the same HyPhy 2.5.100 (MP) binary running RELAX.bf analysis version 4.7.

The nested fits that precede the RELAX mixture also agree, to within two likelihood units: nucleotide GTR log-likelihoods differed by more than 0.01 units in five of eighty pairs and MG94×REV in eighteen of eighty, with medians of 0.0000 and a largest discrepancy of 1.99 (Supplementary S2.1). That largest value sits at the χ^2^_1_ threshold for a one-parameter difference rather than below it, so we report it rather than call the sub-stage fits identical; the RELAX-stage divergences it is being compared against are three orders of magnitude larger. Data, model and software are held fixed and the sub-stage fits recovered; whatever diverges, diverges at the last step alone.

### Parallel arithmetic is not the cause

Message-passing parallelism is excluded by construction, every fit being a single hyphy MP process with HYPHYMPI never invoked. Shared-memory parallelism was excluded by experiment: three genes were each fitted four times under the default thread count and four times under CPU=1, and in all six arms the four replicates returned four distinct values of *K* (Figure 2b) and four distinct log-likelihoods, single-threaded execution widening the spread in two of the three genes. A pre-registered extension repeated the single-threaded condition on twenty null orthologues, the three above and seventeen drawn at random from the remaining eighty-six, two runs each: no gene returned the same log-likelihood twice, the within-gene ranges running from 0.006 to 377 units with a median of 3.2, and one gene fell within 0.01 (Supplementary S2.2).

**Figure 1.**
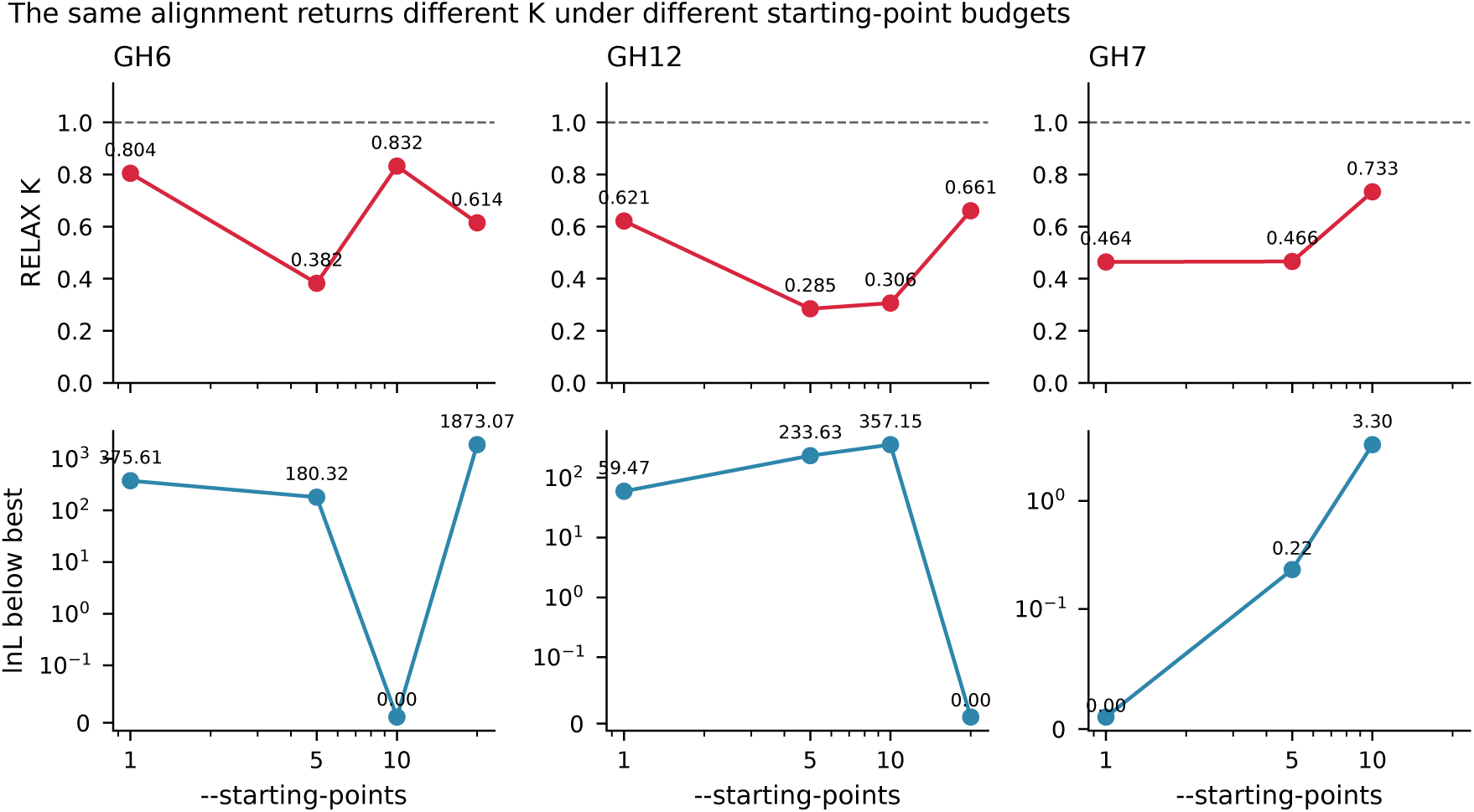
The same alignment returns different fits under different starting-point budgets. One alignment from each of three enzyme families was analysed repeatedly with HyPhy RELAX, changing only the number of optimiser starting points; this is the starting-point arm, extended beyond the one-versus-five comparison to ten and twenty points. Columns are families: GH6, GH12 and GH7. Eleven fits are plotted in all, GH6 and GH12 at 1, 5, 10 and 20 starting points and GH7 at 1, 5 and 10 only, no twenty-point fit being available for GH7. (a) Top row: the RELAX selection-intensity parameter *K* (dimensionless) at each setting, the dashed line marking *K* = 1. *K* takes the values 0.804, 0.382, 0.832 and 0.614 in GH6; 0.621, 0.285, 0.306 and 0.661 in GH12; and 0.464, 0.466 and 0.733 in GH7. (b) Bottom row: the alternative-model log-likelihood of the same fit, expressed in log-likelihood units below the highest value that family returned across the settings shown, on a log scale, so that zero marks the best setting. Those highest values are −52481.56 (GH6, ten starting points), −48169.15 (GH12, twenty) and −104268.37 (GH7, one). **The three bottom panels are scaled independently.** The largest deficit is 1873.07 log-likelihood units in GH6, 357.15 in GH12 and 3.30 in GH7, a 568-fold span, so marker heights are comparable down a column and not across columns. The grid of initial values is drawn at random and was not held fixed, so the panels show variation between settings and not a trend with the number of starting points.

**Figure 2.**
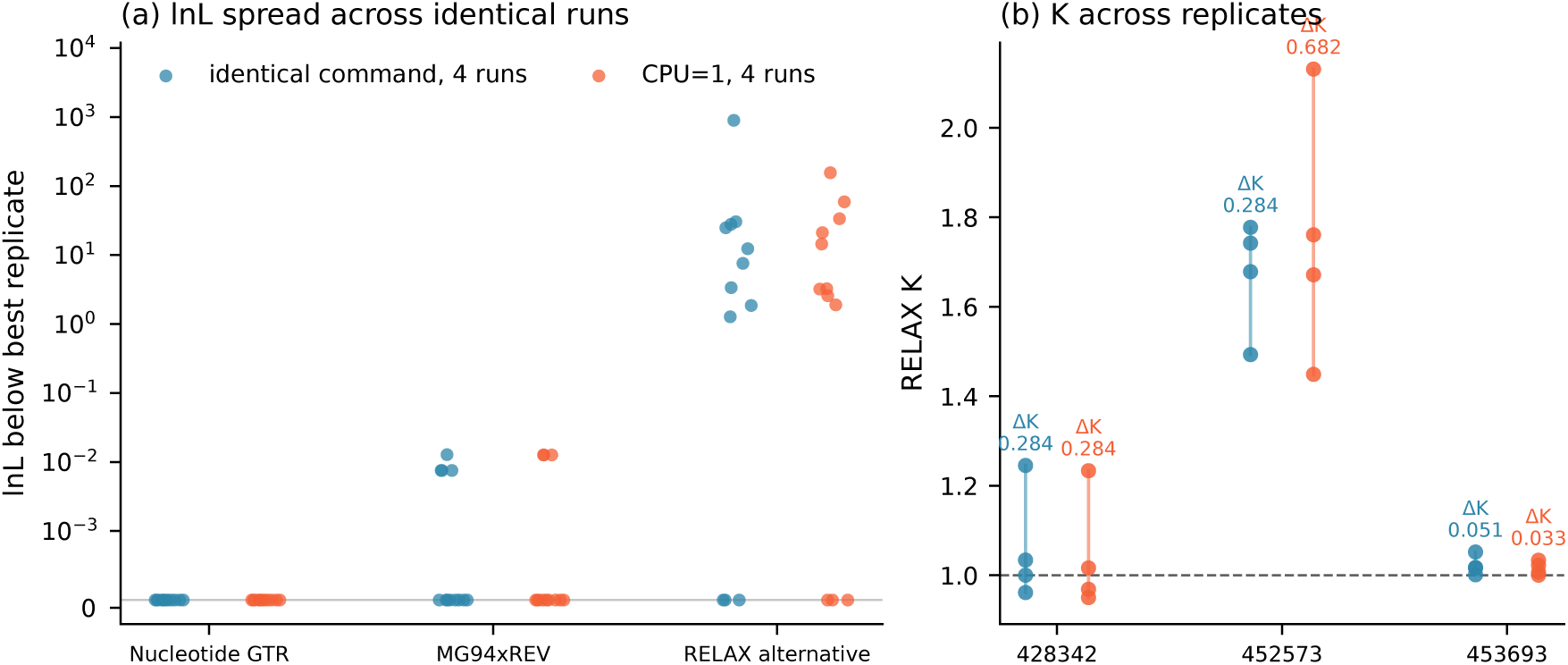
Repeating the same command changes only the last of three nested fits. Log-likelihoods and *K* returned by HyPhy RELAX when a byte-identical command is repeated on the same alignment. Three BUSCO orthologue alignments were each fitted eight times, four replicates under the default thread count and four under single-threaded execution (CPU=1), giving 24 fits in all; the two execution arms are plotted in separate colours throughout. Starting points and branch labelling are not varied here, and the arms differ in thread count alone. (a) For each of the three nested fitting stages, how far below the best replicate each fit falls, in natural-log likelihood units on a logarithmic axis. The nucleotide GTR stage is flat: the largest spread within any gene and arm is 2.7 × 10^-7^, and one arm returns bit-identical values. MG94×REV moves by at most 0.013. The RELAX alternative model, fitted last, moves by 7.6 to 896 units depending on gene and arm. (b) The estimate of *K* across the same replicates, one column per gene, with the within-arm range annotated. It spans 0.033 to 0.682, and for one gene the estimate straddles 1 — the value that separates a relaxation call from an intensification call.

The nucleotide GTR log-likelihood, by contrast, agrees across repeats to within 2.7 × 10^-7^ (Figure 2a), the magnitude expected from a reordered floating-point reduction, and under CPU=1 that residual persists at 8.9 × 10^-8^ and 1.5 × 10^-7^ in two of the three genes; across the twenty single-threaded pairs of the extension the GTR values agreed exactly as printed in fifteen, to within 8.4 × 10^-7^ in four, and differed by 2.5 × 10^-3^ in one, against RELAX-stage differences of 0.006 to 377 in the same pairs. The non-determinism originates above the threading layer, amplified from the eighth significant digit at worst at the GTR stage to tens of likelihood units at the RELAX stage.

### The seed does not control it

RANDOM_SEED is a documented HyPhy variable, assignable on the command line as ENV=expression, but it has no effect on this analysis. Two executions issued with the same seed value returned *K* = 0.290829 with *P* = 1.6 × 10^-4^, a significant relaxation, and *K* = 0.992032 with *P* = 0.887, nothing at all. Setting it through the shell environment failed likewise (Supplementary S2.3). The cause is specific, and it is one the user cannot see. An ENV= assignment is stored as a string and applied after the startup routine has already seeded the generator from the clock and the process identifier; the assignment then sets the variable without calling the seeding routine again. That routine is reached from exactly two places in the source, and the only one a user can reach is the SetParameter path. A batch file that seeds through SetParameter draws reproducibly where the same seed supplied on the command line does not, and in both cases the variable reads back as the value supplied, so nothing in the output distinguishes a seeded run from an unseeded one. We verified this at runtime in 2.5.100, the release under which every analysis here ran, and found the same arrangement in the 2.5.28 sources. Release 2.5.101 revises the startup path to re-seed the generator when a command-line assignment names RANDOM_SEED — the documented form then seeds draws reproducibly — a change its release notes do not mention (Supplementary S2.12).

Reducing the randomness rather than fixing it does not help either: shrinking the Latin hypercube grid of initial values from its default of 250 to 10, the smallest the program accepts, left every completed repeat distinct in both *K* and log-likelihood, with or without a seed (Supplementary S2.4).

Nor is the behaviour particular to the release we used: four repeats of the same command under HyPhy 2.5.28, 2.5.100 and 2.5.101 gave a distinct log-likelihood for every completed repeat in every gene under every version, the current release included (Supplementary S2.5). No documented option makes a RELAX run reproducible under 2.5.100. The analysis can nevertheless be made reproducible, by two measures neither of which suffices alone. Seeding through SetParameter while holding the thread count at one returned identical log-likelihoods, *K* and *P* values in all twenty-one runs of three genes, twelve at one optimiser starting point and nine at five, the output files differing only in their timing fields, and then, in the pre-registered extension, in all sixty runs of the twenty null orthologues, three runs each. Holding the thread count alone left all seven repeat pairs divergent and moved a *P* value across 0.05 in three of them, in one case from 8.0 × 10^-8^ to 0.78, and left all twenty pairs of the extension divergent; seeding alone narrowed the spread of one gene from 7.84 log-likelihood units to 0.048 without closing it (Supplementary S2.12). Under 2.5.101, where the command-line seed does reach the generator, the extension returned the same verdict in the documented form: the seed with the thread count held at one gave one log-likelihood per gene in all sixty runs of the twenty, and the seed at the default thread count reproduced none of five genes in fifteen runs, in one of them returning *K* = 1.0000 exactly with *P* = 1 from one run and *K* = 0.39 with *P* = 0.0013 and 0.0015 from the other two. All four outcomes were as predicted before the runs. What constrains a user is therefore not the absence of a remedy but the absence of a discoverable one: the current release documents the seed, and no documentation we could find states that it binds the fit only when the thread count is held at one.

### The magnitude of the divergence

Repeating an identical command produced divergent fits in every gene examined, two in the seed pilot, three in the threading control and twenty in the single-threaded extension, of at least three kinds: verdict reversal, catastrophic single-run failure, and a stable verdict over an estimate too variable to use (Supplementary S2.6).

Across the eighty null pairs (Supplementary S2.10), not one reproduced its log-likelihood to within 0.01 units: thirty improved with additional starting points and fifty became worse, with a median change of −6.63 and a range from −6520 to +1380. Because the starting grid is drawn at random and was not held fixed, this is not a test of --starting-points, but something weaker and sufficient: the returned fit is not a function of the alignment and the user’s settings alone.

Agreement in *K* is not evidence to the contrary: a parameter pinned at 1.0000 cannot record that the surface underneath it has moved (Results 3).

### The cause is the optimiser, not the data

The remaining explanation is that these alignments are pathological — misaligned, saturated, or poorly resolved. Simulation answers this: there the generating model is exactly the fitted model. The shallow simulated set of 160 replicates described in the companion manuscript (Results 5) was fitted twice under the identical two-arm protocol, at one and at five starting points; 152 returned a statistic under both, and five of those, 3.3 per cent, reproduced their log-likelihood; twenty per cent agreed to within one likelihood unit, the median disagreement was 9.83 units and the largest exceeded 10,000. On data with no alignment error, no rate variation across sites, no multinucleotide substitution and no model misspecification, the optimiser fails to recover its own log-likelihood in roughly ninety-seven per cent of cases. A rate of 0/80 is not distinguishable from 3.3 per cent — the probability of no successes in eighty draws at that rate is 0.069 — so the correct statement is not that the real data are worse, but that clean data are already this bad.

### The instability follows the mixture, not the program

PAML implements models with and without a mixture over sites, so fitting the same alignments from two starting values of *ω* separates behaviour specific to RELAX from behaviour general to codon-model optimisation. Under the two-ratio branch model, which has no site classes, the two starting values returned the same log-likelihood for all 288 genes across the three sets under both null and alternative, the largest difference being 0.0001. Under Clade Model C and M2a_rel, which add three site classes, agreement fell to between 27 and 67 per cent of genes depending on set and model, the largest differences on the null gene set being 8,557 and 6,037 log-likelihood units. Depth compounds this, and in both models: on replicates matched to the divergence of the observed alignments, agreement under M2a_rel falls from 27 and 30 per cent of genes to 5.0, and under Clade Model C from 34 and 40 per cent to 11.2 (Supplementary S2.7). Depth and a mixture over sites together are where this optimiser fails; neither alone produces failure on that scale.

The irreproducibility is therefore neither general to codon-model optimisation nor specific to HyPhy: it is associated with the introduction of a mixture over sites, and appears in two programs with no shared code. A single RELAX analysis of GH15 shows the same transition (Supplementary S2.9).

### More starting points make the calibration worse

Fitting from more starting points was tested on the same simulated alignments under the true null. Among the ninety-four replicates of the shallow five-template set (companion manuscript, Results 5) that converged under both settings, the false-positive rate rose from 17.0 per cent at one starting point to 22.3 per cent at five, while the two visible pathology indicators, terminations with a negative likelihood-ratio statistic and returns of *K* = 1.0000 exactly, both became less frequent (Supplementary S2.8). Additional starting points find a deeper optimum that rejects the null, and suppress the symptoms by which a user might otherwise have noticed the problem.

### Instability is general; its direction is not

The same simulated alignments were re-fitted at the same five starting points with the test label moved from the tips to an internal branch, the two arms differing in the labelled tree and nothing else (companion manuscript, Results 5). The excess rejection rate survives the move in both the shallow and the divergence-matched sets, at 13.5 and 35.0 per cent against a nominal 5: the instability documented here is a property of the optimiser, not of our labelling.

It has also been reported independently by other users. One report of RELAX returning *K* = 2.14 and then *K* = 0.68 on an identical rerun, the verdict crossing from intensification to relaxation, and another of the same non-determinism in aBSREL, were opened against the HyPhy repository in 2024 and closed by an automated stale-bot without resolution (veg/hyphy issues 1685 and 1722, accessed 20 August 2026).

## RESULTS 2 — THE SOFTWARE’S OWN DIAGNOSTICS, AT DEFAULT SETTINGS

RELAX carries convergence diagnostics that are on by default and writes them into its own output. Of the 410 fits in the observed analysis and its null distributions — 350 on null orthologues and 60 on the enzyme entries, 175 of them under a randomised test set and 235 under the real ectomycorrhizal labelling — 306 (74.6 per cent) carry at least one such flag. The figure is a census of the fits this study performed, pooled across arms, not an estimate of a rate (Supplementary S2.11). Two of these diagnostics describe states whose consequences cannot be inferred from the published output.

### Terminations that are impossible under convergence

When RELAX obtains a negative likelihood-ratio statistic it re-fits the alternative/null pair from a fresh starting point, up to five times. In 46 of 410 fits (11.2 per cent) the five rescue re-fits were exhausted and the analysis terminated with the statistic still negative. That proportion pools fits made at one, five and ten optimiser starting points, and the arms differ; among the ectomycorrhizal-free null genes alone (null C) the figure is 15 of 81 (18.5 per cent).

A negative likelihood-ratio statistic means the unconstrained alternative model was fitted to a worse likelihood than the null model nested inside it. If either fit had converged, this could not occur: the null is a point in the alternative’s parameter space. These 46 fits are not imprecise; they are demonstrably not at a maximum, and the software reports them with a *P* value like any other. The observation depends on no control, no replicate, no seed and no simulation.

### The reported K is often not the unconstrained estimate

Before finalising a fit, RELAX profiles *K* over a 200-point grid from 0 to 4.975 (Figure 3) and compares the best likelihood on either side of *K* = 1. Where those two maxima lie within five likelihood units — 184 fits, 44.9 per cent — it declares the surface flat and re-fits with *K* restricted to the opposite side of one. In 123 fits (30.0 per cent; 37.0 per cent of the ectomycorrhizal-free nulls) that restricted re-fit achieved a higher likelihood and its value replaced the original. The reported *K* is then not the unconstrained maximum-likelihood estimate but a re-estimate under a domain restriction, and the output table does not distinguish the two. Any downstream comparison of *K* values, including the one in our own earlier analysis, therefore combines quantities of two different kinds.

**Figure 3.**
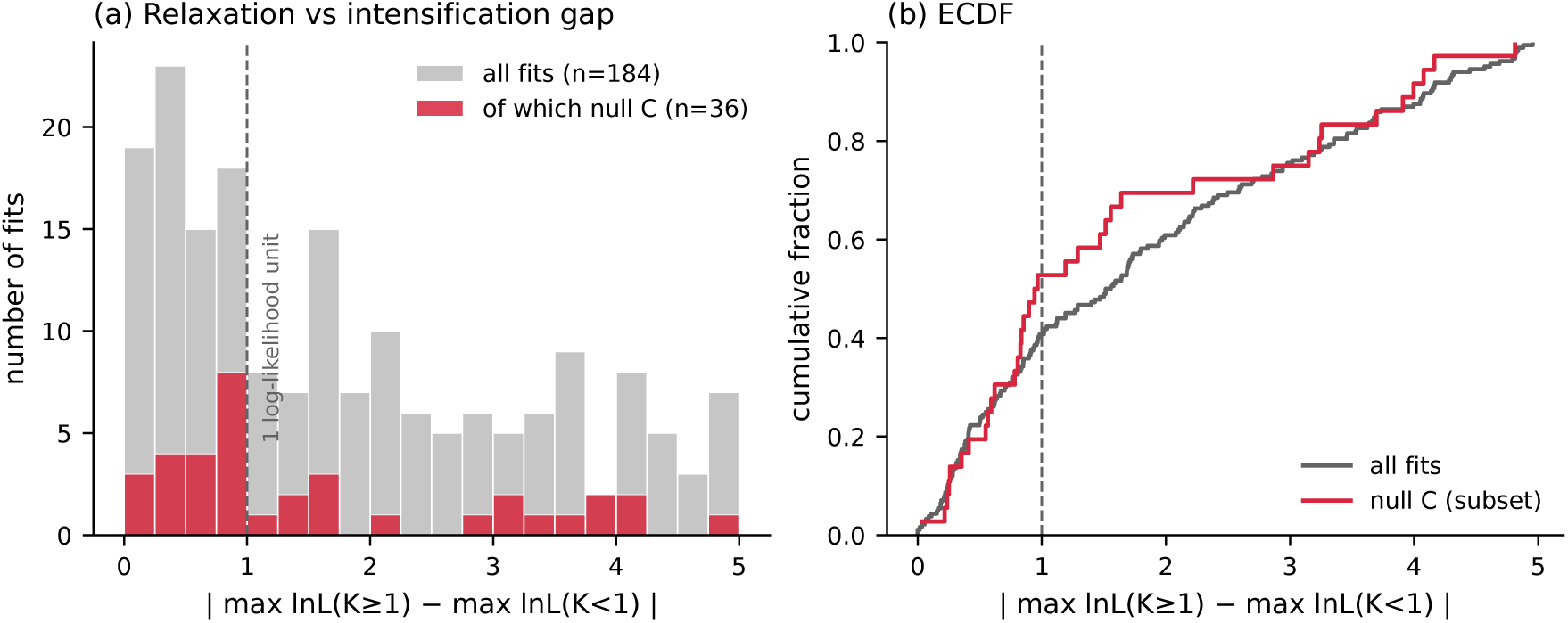
RELAX’s own flat-surface diagnostic. Before finalising a fit, RELAX compares the best log-likelihood reachable with *K* >= 1 against the best reachable with *K* < 1, flagging the fit when the two lie within five log-likelihood units of each other. Plotted is the absolute difference between those two maxima, in log-likelihood units, for every fit in which the check fired: 184 fits, pooled over the eight arms in which it fired at all. By gene, 162 are null orthologues and 22 are the enzyme entries themselves. By labelling, 87 come from a randomised test set — 47 from the contaminated random-tip null (A), 36 from the ectomycorrhizal-free null (C) and 4 from the ten-starting-point check — and 97 from the real ectomycorrhizal labelling: 75 on null orthologues (null B at one, five and ten starting points) and 22 on the enzyme entries. **The pooled histogram is therefore not a null distribution**; the null C subset is drawn separately. (a) Histogram of the difference, number of fits per bin, all 184 fits in grey with the 36 null C fits overlaid in red. (b) Empirical cumulative distribution of the same quantity, all fits and the null C subset drawn separately. The dashed line in both panels marks one log-likelihood unit. Medians are 1.54 units over all 184 fits and 0.95 over the 36 null C fits. The distribution is censored at five units by the rule that selects fits into it, so the maximum of 4.96 is a property of that rule rather than a measurement; the medians, which lie well inside the bound, are not.

The profile also quantifies how weak the distinction is, for the fits in which it fired. Among those 184 the median separation between the best relaxed and the best intensified solution is 1.54 likelihood units, and 0.95 among the 36 of 81 ectomycorrhizal-free nulls in which it fired.

## RESULTS 3 — THE ENZYME ENTRIES: ONE FAILED RUN, AND TWENTY-SEVEN PAIRS THAT DO NOT REPRODUCE

### The tested entries and the resolution available

Twenty-eight plant-cell-wall-degrading enzyme entries had codon alignments deep enough to test: twenty families, seven *GH5* subfamilies and a subsampled *GH5*. Twenty-seven returned a fitted *K* under both starting-point settings.

The twenty-eighth, *GH5_22*, did not, and what happened to it is itself a result. Its five-starting-point run terminated on the optimiser’s own consistency check, which reported that the log-likelihood at the final point did not match the best the run had recorded, 106.82 units apart, and it left an empty output file that the batch did not re-attempt. Re-running that same command three times returned a fit on every attempt, and three different ones: *K* from 0.900 to 1.096, crossing the null value so that the direction of the inferred effect reverses; log-likelihoods 113.74 units apart; and *P* from 0.038 to 0.515, reaching the conventional threshold in one run of the three, where the entry would have been reported as relaxed. We report it as a failure rather than dropping it, and do not add it to the paired set, since no single five-starting-point value exists to be paired.

### The entries do not reproduce

Table 1 lists all twenty-seven paired entries ordered by the change in log-likelihood between the two settings, not by *K*; sorting by *K* would present that column as an effect size. Not one entry reproduced its log-likelihood to within 0.01 units; the median absolute change was 103 units and the largest exceeded 3,400. Two entries, *GH10* and *GH43*, returned *K* identical to six decimal places, in both cases exactly 1.0000, while their log-likelihoods moved by 581 and 607 units respectively. Both would pass a *K*-based reproducibility check, and would pass it because *K* was 1.0000 in both fits while the likelihood moved by hundreds of units (Results 1).

**Table 1.** Every enzyme entry refitted from a different optimiser starting configuration, ordered by how far its log-likelihood moved. Each entry was fitted twice with RELAX under commands differing only in the number of optimiser starting points, one against five; alignment, tree and branch partition were identical in every pair (Results 1). *K* is reported as evidence of instability and not as an estimate (Discussion 2). Flags abbreviate the convergence diagnostics RELAX writes into its own output for the five-starting-point fit, and the strings to grep for are convergence-negative-lrt, convergence-flat-surface and convergence-unstable. Here neg-lrt(p*n*) records a negative likelihood-ratio statistic surviving *n* rescue passes and LRT < 0 exhausted one that survived all five, flat the flat-surface check with the likelihood gap it reported, and K-replaced a fit whose *K* the program re-estimated under a domain restriction (Results 2). GH7, the entry discussed in Discussion 2 and the one that drops out of significance under the matched null (companion manuscript, Results 3), is the last row: its two fits agree to 0.22 likelihood units, and a third configuration disagreed with both. Twenty-eight entries were attempted; GH5_22 produced no output under the five-starting-point configuration, its run having terminated on the optimiser’s internal consistency check, and is absent, leaving the 27 paired entries above (what it returns on re-running is given in Results 3). Across them the median |ΔlnL| is 102.87, with a range of 0.22 to 3474.08. The median |Δ*K*| is 0.0314 with a range of 0.0000 to 0.5947, a figure that pools eight entries whose reported *K* is a re-estimate under a domain restriction with nineteen whose *K* is an unconstrained maximum (Results 2). **No entry reproduced its log-likelihood to within 0.01 units.** Fifteen entries reached *P* < 0.05 under each configuration, but not the same fifteen: four reversed their call between the two, CE1, GH5, CE12 and GH5_9. The only two entries whose *K* matched exactly, GH43 and GH10, both returned *K* = 1.0000 exactly while their log-likelihoods differ by 606.85 and 581.50 units — agreement in the parameter recording that it could not move rather than that the fit had converged (Discussion 2). Where a *P* value is printed as < 10^-16^ the raw value underflowed double precision and we do not report a figure for it.

| Entry | Se<br>qs | Sit<br>es | $K$<br>(sp<br>1) | $p$ (sp<br>1) | $\ln L$ (sp<br>1) | $K$<br>(sp<br>5) | $p$ (sp<br>5) | $\ln L$ (sp<br>5) | $\vert \Delta \ln L\vert $ | $\vert \Delta K\vert $ | Flags<br>(sp 5) |
| --- | --- | --- | --- | --- | --- | --- | --- | --- | --- | --- | --- |
| CE8 | 134 | 331 | 0.75<br>13 | $1.2 \times 10^{-6}$ | -<br>85,003.<br>97 | 1.34<br>60 | $< 10^{-16}$ | -<br>88,478.<br>05 | 3474.<br>08 | 0.59<br>47 | — |
| GH5_<br>15 | 80 | 463 | 0.55<br>30 | $< 10^{-16}$ | -<br>71,241.<br>10 | 0.65<br>07 | $5.9 \times 10^{-8}$ | -<br>74,319.<br>55 | 3078.<br>46 | 0.09<br>77 | — |
| CE1 | 161 | 275 | 0.77<br>38 | $6.5 \times 10^{-5}$ | -<br>79,725.<br>32 | 1.00<br>84 | 0.71<br>3 | -<br>82,003.<br>31 | 2277.<br>99 | 0.23<br>46 | neg-lrt(p5),<br>flat<br>3.74,<br>K-replace<br>d |
| GH5 | 902 | 353 | 1.00<br>00 | 1 | -<br>474,429<br>.82 | 0.96<br>86 | $2.4 \times 10^{-9}$ | -<br>475,151<br>.98 | 722.1<br>6 | 0.03<br>14 | neg-lrt(p1) |
| CE16 | 279 | 288 | 0.87<br>47 | 0.00<br>19 | -<br>159,075<br>.89 | 0.89<br>42 | 0.00<br>40 | -<br>158,425<br>.20 | 650.6<br>9 | 0.01<br>95 | neg-lrt(p1) |
| GH43 | 331 | 297 | 1.00<br>00 | 1 | -<br>175,020<br>.15 | 1.00<br>00 | 1 | -<br>174,413<br>.30 | 606.8<br>5 | 0.00<br>00 | neg-lrt(p3),<br>flat<br>2.18,<br>K-replace<br>d |
| GH10 | 164 | 355 | 1.00<br>00 | 1 | -<br>116,311<br>.09 | 1.00<br>00 | 1 | -<br>116,892<br>.58 | 581.5<br>0 | 0.00<br>00 | neg-lrt(p3),<br>flat<br>1.43,<br>K-replace<br>d |
| GH51 | 94 | 643 | 0.77<br>45 | 0.02<br>37 | -<br>120,825<br>.35 | 0.77<br>32 | $7.4 \times 10^{-7}$ | -<br>120,316<br>.22 | 509.1<br>3 | 0.00<br>13 | neg-lrt(p1) |
| CE12 | 73 | 259 | 0.74<br>60 | 0.02<br>78 | -<br>40,816.<br>45 | 1.00<br>00 | 1 | -<br>41,227.<br>63 | 411.1<br>8 | 0.25<br>40 | LRT <<br>0<br>exhaust ed, flat<br>1.70,<br>K-replace<br>d |
| GH5_<br>9 | 279 | 470 | 0.98<br>41 | 0.14<br>2 | -<br>240,512<br>.60 | 0.82<br>19 | $< 10^{-16}$ | -<br>240,125<br>.26 | 387.3<br>5 | 0.16<br>22 | neg-lrt(p1) |
| GH5_<br>12 | 93 | 752 | 0.85<br>21 | $5.9 \times 10^{-8}$ | -<br>140,885<br>.86 | 0.88<br>40 | $4.7 \times 10^{-8}$ | -<br>141,107<br>.89 | 222.0<br>2 | 0.03<br>19 | — |
| GH6 | 69 | 397 | 0.80<br>45 | $1.8 \times 10^{-4}$ | -<br>52,857.<br>18 | 0.38<br>24 | $4.8 \times 10^{-9}$ | -<br>52,661.<br>88 | 195.3<br>0 | 0.42<br>21 | neg-lrt(p1) |
| GH12 | 92 | 251 | 0.62<br>13 | $6.7 \times 10^{-9}$ | -<br>48,228.<br>61 | 0.28<br>46 | $< 10^{-16}$ | -<br>48,402.<br>78 | 174.1<br>6 | 0.33<br>67 | — |
| CE15 | 48 | 396 | 1.21<br>66 | 0.06<br>29 | -<br>39,437.<br>76 | 1.20<br>14 | 0.05<br>7 | -<br>39,540.<br>63 | 102.8<br>7 | 0.01<br>52 | neg-lrt(p1),<br>flat<br>4.82,<br>K-replace<br>d |
| PL4 | 60 | 525 | 1.00<br>01 | 1 | -<br>69,252.<br>42 | 1.00<br>85 | 0.77<br>3 | -<br>69,348.<br>99 | 96.57 | 0.00<br>84 | neg-lrt(p1),<br>flat<br>3.20 |
| GH5_<br>50 | 43 | 509 | 0.80<br>89 | $7.6 \times 10^{-4}$ | -<br>51,791.<br>62 | 0.76<br>52 | 0.00<br>81 | -<br>51,853.<br>90 | 62.27 | 0.04<br>37 | neg-lrt(p1) |
| CE5 | 162 | 212 | 0.91<br>33 | 0.00<br>48 | -<br>69,492.<br>25 | 0.83<br>57 | 0.00<br>26 | -<br>69,461.<br>69 | 30.56 | 0.07<br>76 | neg-lrt(p1) |
| GH5_<br>5 | 159 | 383 | 1.00<br>00 | 1 | -<br>99,763.<br>74 | 1.02<br>63 | 0.83<br>7 | -<br>99,790.<br>71 | 26.97 | 0.02<br>63 | neg-lrt(p4),<br>flat<br>0.41,<br>K-replace<br>d |
| GH5_<br>7 | 91 | 392 | 1.02<br>43 | 0.93<br>7 | -<br>69,891.<br>58 | 1.00<br>47 | 0.93<br>0 | -<br>69,918.<br>40 | 26.81 | 0.01<br>96 | neg-lrt(p2),<br>flat<br>0.15,<br>K- replace<br>d |
| GH74 | 45 | 746 | 0.84<br>18 | 0.55<br>7 | -<br>61,546.<br>14 | 0.89<br>04 | 0.24<br>2 | -<br>61,524.<br>01 | 22.13 | 0.04<br>86 | flat -<br>2.08 |
| GH53 | 46 | 351 | 0.61<br>63 | 0.00<br>25 | -<br>34,344.<br>36 | 0.66<br>75 | 0.01<br>1 | -<br>34,328.<br>49 | 15.87 | 0.05<br>11 | — |
| GH5.s<br>ub | 160 | 353 | 0.96<br>09 | 0.13<br>8 | -<br>110,452<br>.88 | 0.99<br>20 | 0.47<br>6 | -<br>110,439<br>.35 | 13.54 | 0.03<br>11 | neg-lrt(p4),<br>flat<br>0.50,<br>K-replace<br>d |
| GH28 | 289 | 376 | 0.86<br>86 | $2.7 \times 10^{-5}$ | -<br>200,469<br>.89 | 0.86<br>94 | $2.6 \times 10^{-5}$ | -<br>200,462<br>.96 | 6.93 | 0.00<br>07 | neg-lrt(p1) |
| GH93 | 46 | 376 | 0.90<br>44 | 0.47<br>2 | -<br>38,600.<br>95 | 0.87<br>81 | 0.27<br>2 | -<br>38,597.<br>27 | 3.68 | 0.02<br>63 | — |
| PL1 | 118 | 302 | 0.87<br>69 | 0.10<br>3 | -<br>74,292.<br>26 | 0.87<br>77 | 0.09<br>23 | -<br>74,295.<br>67 | 3.41 | 0.00<br>08 | neg-lrt(p1) |
| PL3 | 84 | 251 | 0.55<br>52 | $6.9 \times 10^{-4}$ | -<br>30,994.<br>60 | 0.63<br>83 | $2.9 \times 10^{-6}$ | -<br>30,996.<br>24 | 1.64 | 0.08<br>31 | — |
| GH7 | 140 | 507 | 0.46<br>42 | $1.6 \times 10^{-9}$ | -<br>104,268<br>.37 | 0.46<br>60 | $2.3 \times 10^{-9}$ | -<br>104,268<br>.59 | 0.22 | 0.00<br>19 | neg-lrt(p1) |

Four entries changed their call between the two settings: *CE1*, *CE12*, *GH5* and *GH5_9*. Fifteen of twenty-seven were significant at one starting point and fifteen at five, but not the same fifteen; reporting only the count would have found the two settings in perfect agreement.

The diagnostics of Results 2 fall unevenly across the table: nineteen of the twenty-seven entries carry a negative likelihood-ratio flag, ten triggered the flat-surface check, and in eight the reported *K* is a re-estimate under a domain restriction rather than the unconstrained maximum. Those eight include two *GH5* subfamilies and the subsampled *GH5* alignment, on which our earlier account of subfamily-level divergence rested.

## DISCUSSION 1 — FAILURES THAT DO NOT ANNOUNCE THEMSELVES

These failures share a form, and it is the form rather than any one of them that we would put to a reader. In each case the step terminated normally and returned a value of the kind we were looking for, and nothing in its output separated it from a step that had worked. The form is not confined to one program, nor even to the programs. Two of the entries below were observed in two codebases that share none of each other’s, and the first was already present in the database field from which the analysis took its labels.

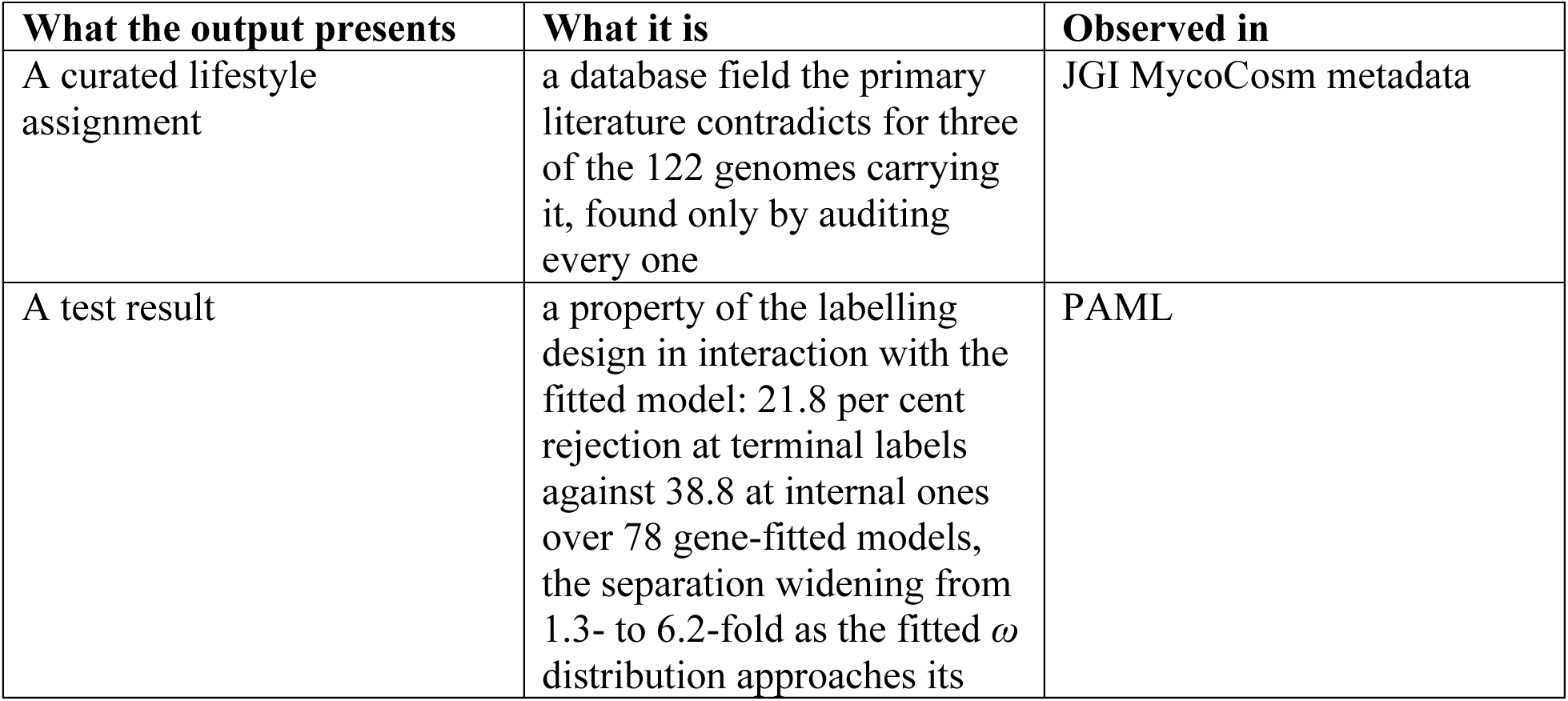
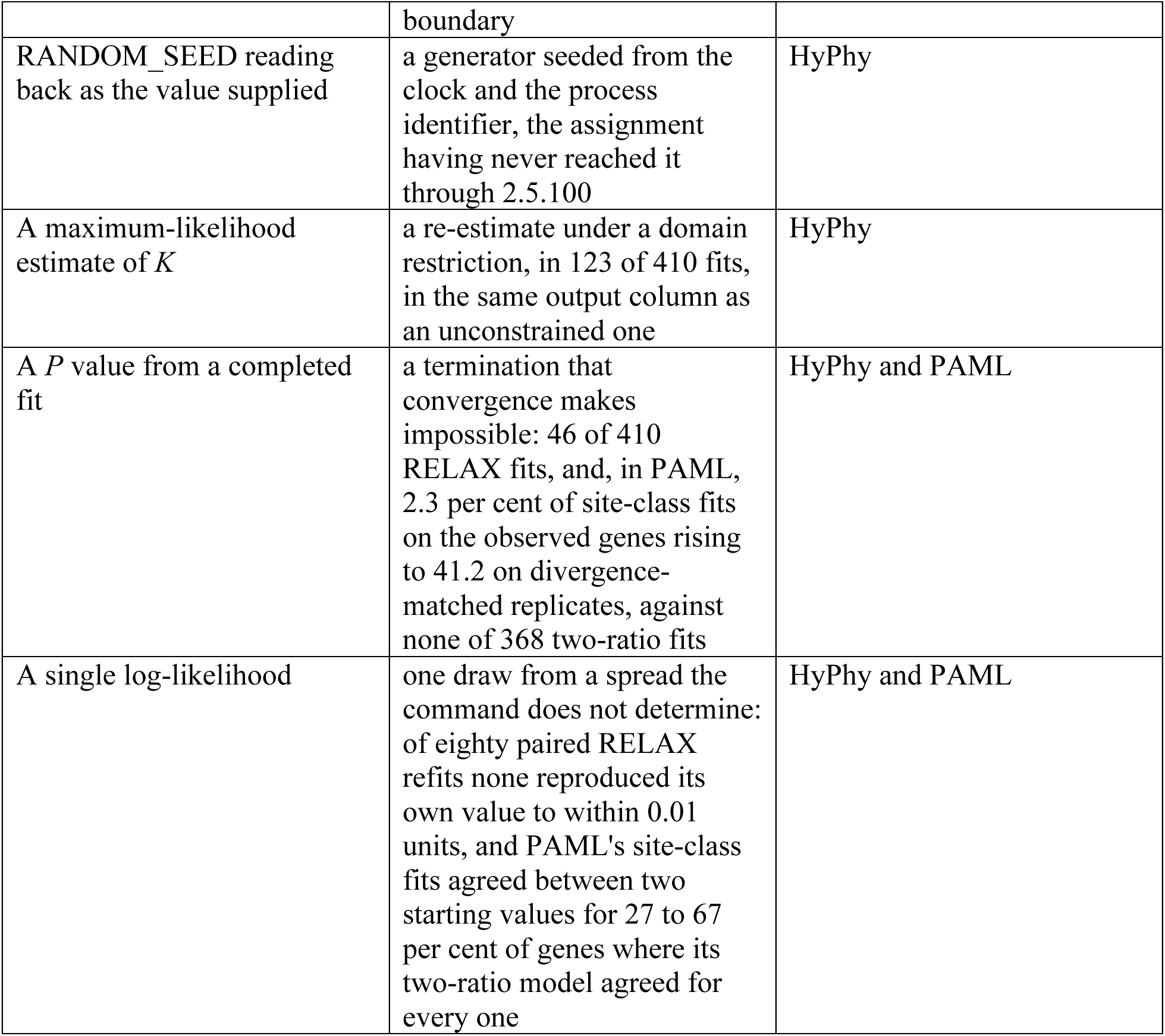

Every method fails somewhere, and a method that announced its failures would need no calibration exercise. What these six have in common is that they do not announce them, and that each is discoverable only by an experiment the output gives no reason to perform. The first is the only one we caught before it reached a result, and it marks the sense in which the count and the selection intensity are not two results of differing strength but two results of differing kind: one can be checked by counting again, and the other cannot be checked by reading the output at all. If these failures are discoverable only by an experiment the output gives no reason to perform, the literature should show few people performing it, and it does. Of 234 papers we examined that use RELAX, none reported the random-number seed of the analysis; two reported the number of optimiser starting points, and between three and five described repeating a fit to check that it reproduced (Supplementary S2.13). Nothing in any output told those authors a seed mattered, and by the mechanism set out in Results 1 a seed recorded in the documented form would not have bound the generator in any case. A reporting standard cannot be met by authors whose tools do not show them what there is to report.

## DISCUSSION 2 — WHAT TO DO INSTEAD

The failures documented here arise from a widely used implementation at its default settings, on data of ordinary size, in a design that comparative genomics increasingly favours. We state what we would now require of any branch-partitioned selection test, tying each recommendation to the observation that forces it.

### Report the log-likelihood of every fit alongside the parameter it produced

A parameter that agrees between two runs is not evidence that either run converged. The only two enzyme entries whose *K* matched to six decimal places were also the two whose log-likelihoods differed by 581 and 607 units. The agreement existed because *K* was at 1.0000 in both fits, where it is uninformative about the likelihood surface.

### Treat agreement between two configurations as insufficient. Fit from at least three, take the best likelihood, and report the spread

One family returned *K* = 0.4642 at one starting point and *K* = 0.4660 at five, the log-likelihoods 0.22 units apart and the test significant at both settings; at ten starting points the same alignment returned *K* = 0.7334 and a *P* value of 0.29. The parameter had moved by 0.27 and the verdict had reversed, at a cost of 3.3 log-likelihood units. Three configurations are a floor rather than a target. In other families the best of four fits returned no significant result (Supplementary S3.1).

### Do not report a point estimate of the parameter of interest; report a profile interval

On simulated data fitted from five starting points, where the true value was 0.7 our recovered estimates ranged from 0.014 to 1.025; where the true value was 1.0, from 0.411 to 2.411. That spread is wider than the range of effect sizes the observational analysis was attempting to distinguish.

### Treat a returned value of exactly 1.000 with particular suspicion

The null value is where a failed optimisation is least visible, since a parameter that has not moved off it looks like a converged estimate of no effect. Among our null genes, 22 per cent returned *K* = 1.0000 exactly, and the family-level result we relied on most heavily in earlier work returned *K* = 1.0000 with a likelihood-ratio statistic of zero, whose likelihood moved by 722 units on re-fitting.

### Record the numerical environment, including the thread configuration

In a phylogenetic generalised least squares model unrelated to the tests above, whether the optimiser converged or terminated abnormally depended deterministically on the number of BLAS threads: ten replicates per condition gave ten successes at eight threads and ten failures at one and at seventy-two, a dependence that is not monotone and that nothing in the output identifies. Where the analysis draws on a random number generator, fix that too, and check that the mechanism you use actually reaches it; in the program tested here the documented form did not reach it in the release we used, and in the current release reaches it but binds the fit only with the thread count held at one (Results 1). The test of whether an environment is pinned is cheap and no substitute exists for it. Run the same command twice and compare the log-likelihoods.

A route to reproducibility exists, then, but not one the documentation leads to; we found it by reading the source, and confirmed it on twenty genes under two releases. The current release documents the seed; that the seed binds the fit only with the thread count held at one, which it did in sixty of sixty runs and failed to do for any of five genes at the default thread count, is documented nowhere we could find. Until it is documented, and better still until it is the default, an author cannot reasonably be expected to have used it, and a reader cannot tell from a methods section whether it was used. What can be asked in the meantime is weaker and still worth asking. It is not that a seed be recorded, since a recorded seed need not bind, but that the fit be repeated and the dispersion published, which here is large enough to change conclusions.

## DISCUSSION 3 — LIMITATIONS

### The nested fits agree to within two likelihood units, not exactly

The nucleotide and codon fits used to exclude data and model artefacts (Results 1) agree to within two likelihood units rather than exactly, and we report that rather than call them identical.

### The evidence concerns one program family, and one release series of it

Every RELAX fit here ran under HyPhy 2.5.100, with repeats under 2.5.28 and 2.5.101; the second implementation, PAML, was used to localise the instability and not to characterise its own. Other branch-partitioned tests in HyPhy, aBSREL and BUSTED among them, were not examined.

### The controls are small

The threading experiment used three genes and the seed pilot two; the pre-registered extension took the single-thread and seeded arms to twenty null orthologues and the seeded default-thread arm to five, all single-copy genes of ordinary size at one starting point, so no enzyme entry and no deeper alignment has been through it. The eighty paired refits vary the number of starting points without holding the starting grid fixed, so they show that the returned fit is not a function of the alignment and the settings alone, and not how it depends on either; the reduced-grid experiment went to the smallest grid the program accepts and no further.

### The simulated evidence is from the shallow set

The 3.3 per cent reproducibility rate was measured on 152 replicates about an order of magnitude less diverged than the observed alignments. On the divergence-matched set it was the site-class models of PAML, not RELAX, that were refitted from two starting values, so the depth dependence reported for those models has no RELAX counterpart here.

### The literature survey counts what is stated

A study that seeded its run without saying so, or repeated a fit and reported only the final one, is counted as not having done so (Supplementary S2.13).

## Supporting information

Supplementary Information (Methods S1, detail S2-S3, Table S4)

## ACKNOWLEDGEMENTS

A portion of these data were produced by the US Department of Energy Joint Genome Institute (https://ror.org/04xm1d337; operated under Contract No. DE-AC02-05CH11231) in collaboration with the user community. A portion of this research was performed under the Facilities Integrating Collaborations for User Science (FICUS) program (proposals: 10.46936/10.25585/60008430 and 10.46936/10.25585/60008431) and used resources at the DOE Joint Genome Institute (JGI) (https://ror.org/04xm1d337) and the National Energy Research Scientific Computing Center (NERSC) (https://ror.org/05v3mvq14), which are DOE Office of Science User Facilities operated under Contract No. DE-AC02-05CH11231, and we thank David Catcheside for permission to include the one use-restricted genome in the panel.

## DATA AVAILABILITY

All data and code underlying this analysis are deposited at Zenodo (doi:10.5281/zenodo.22725503; all versions: doi:10.5281/zenodo.22192658; the version described here is 1.3); the 78-gene simulation panel regenerates from the recorded seeds included there.

The deposit is 1,776 files and 285 MB in version 1.3, assembled by scripts/build_deposit.py, whose MANIFEST pairs each canonical result file with the script that regenerates it. It contains 157 codon alignments, deposited both untrimmed and trimmed as 313 files, restricted to the 35 published genomes among the 51 with coding sequence available: the trimmed set retains 9,620 sequences, and those belonging to the 16 genomes unpublished at the time of analysis are withheld pending the replies described below. Trees carry no sequence and are deposited over all 51 tips — 158 gene trees and 118 labelled trees for the analysed entries and the orthologue null genes, the 88 null C labelled trees, 40 for the enzyme-family control and five for the internal labelling — as are the 304 simulated alignments that no seed can reproduce, held in three subdirectories by the set they belong to; the 312 of the 78-gene panel regenerate instead, as above. For the count analyses the deposit carries the per-genome enzyme count table, the 182-tip species tree, the lifestyle assignments before and after the annotation audit, and the fitting scripts, which together regenerate every estimate of the companion study (Results 2). It further contains the 38 canonical summary result files behind the figures, tables and reported figures of this manuscript and its companion, the six figure audit tables recording every plotted value at full precision, and 170 analysis scripts.

### Two classes of artefact cannot be regenerated from a seed, and are deposited for that reason rather than for convenience

For null C, the routine that drew the random test sets combined its seed constant with Python’s built-in hash() of the gene name, which is salted per interpreter process, so re-running that script does not reproduce the draws that were analysed; the 88 null C trees and the five internal-labelling trees define those analyses and are deposited individually. Draws for the enzyme-family control and for the test-set-size ladder key a CRC32 checksum of the family or gene name instead and do reproduce exactly from the seeds recorded in Methods; they are deposited nonetheless.

The simulated alignments are the stronger case. The generator passes no seed to the sequence simulator, which therefore drew from an unseeded stream, so neither we nor anyone else can reproduce the replicates that were analysed. All 304 are deposited individually, 160 shallow and 144 divergence-matched, together with the per-replicate record of template gene, generating *K*, branch-length scale and realised pairwise identity. They define the reproducibility rate reported in Results 1, and the simulation results of the companion manuscript (Results 5). We describe both defects rather than silently depositing the outputs, because they are instances of the class of problem this paper is about.

Genome assemblies, predicted proteomes and coding sequences were obtained from the JGI MycoCosm portal and from NCBI and are not redistributed here. Portal identifiers for all 51 genomes are listed in Supplementary Table S4. Of these, 16 were unpublished at the time of analysis, meaning that their portal pages carried no associated publication. Consulting the portal’s own use-restriction flag on 31 August 2026 identified one of the sixteen, *Amanita* aff. *grandis* KIS10, as use-restricted; the other fifteen were not so flagged. That genome was sequenced under JGI proposal 1956 (award doi:10.46936/10.25585/60001059, "Acquisition of the sequestrate (truffle like) habit by basidiomycete macrofungi"), and it is used here with the permission of the proposal’s principal investigator, David Catcheside, who asked that the sequencing project be cited: the high-molecular-weight DNA extraction methodology it developed is described in Burgoyne *et al*. (2025), and the genomes it produced are described in Nilsen *et al*. (*Communications Biology*, pending). Those genomes are identified in Supplementary Table S4 and readers should obtain them from the portal directly. The count-based analyses do not depend on the unpublished genomes: restricting the primary contrast to public genomes alone gives 0.411 (0.316–0.535). The selection analyses do depend on them, since all sixteen carry coding sequence and lie inside the 51-genome codon panel, whose other 35 members are published.

For the reproducibility analyses the deposit carries the paired refits of the 27 entries and of the 80 null orthologues, the convergence-flag census and its provenance table, the repeats across HyPhy releases, and the literature-survey script with its record. It further carries the seeded RELAX batch file and the few-second seed demonstration, the raw HyPhy outputs of the seed, thread, mixture-dimension and pre-registered twenty-gene experiments, and the GH5_22 re-executions, added in version 1.3 of the deposit (all versions: doi:10.5281/zenodo.22192658).

Software versions and the release of each annotation database are given in Methods. No new sequence data were generated.

## AUTHOR CONTRIBUTIONS

M.K. designed the analyses, performed the selection analyses and the reproducibility experiments, wrote the simulation and audit code, and drafted the manuscript. J.-H.S. supervised the work and revised the manuscript. Both authors read and approved the final version.

## REFERENCES

Burgoyne LA, Nilsen AR, Lebel T, Catcheside PS, May TW, Orlovich D, Kuo A, Lipzen A, LaButti K, Riley R, Andreopoulos W, Koriabine M, Yan M, Ng V, Grigoriev IV, Catcheside DEA. 2025. Methodology for extracting high-molecular-weight DNA from field collections of macrofungi. J Fungi. 11:490.

HyPhy issue tracker. 2024. Issue 1685, RELAX empty files in results, and results differing when re-run, opened 30 January 2024, closed 1 May 2024 by an automated stale-bot; issue 1722, Different aBSREL results for same inputs, opened 14 July 2024, closed 29 September 2024 likewise. github.com/veg/hyphy/issues (accessed 20 August 2026).

Kim M, Shin J-H. 2026. Relaxed selection and convergent loss erode the retained plant-cell-wall-degrading enzymes of ectomycorrhizal fungi. bioRxiv 2026.07.20.739482, version 1. doi:10.64898/2026.07.20.739482.

Kosakovsky Pond SL, Poon AFY, Velazquez R, Weaver S, Hepler NL, Murrell B, Shank SD, Magalis BR, Bouvier D, Nekrutenko A, Wisotsky S, Spielman SJ, Frost SDW, Muse SV. 2020. HyPhy 2.5—a customizable platform for evolutionary hypothesis testing using phylogenies. Mol Biol Evol. 37:295–299.

Kosakovsky Pond SL, Wisotsky SR, Escalante A, Magalis BR, Weaver S. 2021. Contrast-FEL—a test for differences in selective pressures at individual sites among clades and sets of branches. Mol Biol Evol. 38:1184–1198.

Manni M, Berkeley MR, Seppey M, Simão FA, Zdobnov EM. 2021. BUSCO update: novel and streamlined workflows along with broader and deeper phylogenetic coverage for scoring of eukaryotic, prokaryotic, and viral genomes. Mol Biol Evol. 38:4647–4654.

Muse SV, Gaut BS. 1994. A likelihood approach for comparing synonymous and nonsynonymous nucleotide substitution rates, with application to the chloroplast genome. Mol Biol Evol. 11:715–724.

Nilsen AR, Chyou D, Plett JM, Dobbie F, Jackson C, McBride TM, Daum C, Yoshinaga Y, Lipzen A, LaButti K, Riley RW, Andreopoulos W, Kuo A, Ng V, Grigoriev IV, Burgoyne LA, Catcheside PS, Lebel T, May TW, Brown CM, Catcheside DEA, Orlovich DA. Divergent genetic routes to convergent sporing body forms in truffle-like fungi. Communications Biology, pending.

Spielman SJ, Wilke CO. 2015. Pyvolve: a flexible Python module for simulating sequences along phylogenies. PLoS One 10(9):e0139047.

Weadick CJ, Chang BSW. 2012. An improved likelihood ratio test for detecting site-specific functional divergence among clades of protein-coding genes. Mol Biol Evol. 29:1297–1300.

Wertheim JO, Murrell B, Smith MD, Kosakovsky Pond SL, Scheffler K. 2015. RELAX: detecting relaxed selection in a phylogenetic framework. Mol Biol Evol. 32:820–832.

Yang Z. 2007. PAML 4: phylogenetic analysis by maximum likelihood. Mol Biol Evol. 24:1586–1591.

