## Supplementary Information (Methods S1, detail S2-S3, Table S4) for "RELAX does not reproduce its own estimates at default settings, and its output does not show it"

### SUPPLEMENTARY METHODS S1 — PROCEDURAL DETAIL FOR THE METHODS (RELAX INVOCATION, REPRODUCIBILITY, NULL DISTRIBUTIONS, SIMULATION)

*Supports M2 (RELAX invocation)*

The canonical invocation was

```
hyphy relax --alignment <codon.fasta> --tree <labelled.nwk>  --test Test --
starting-points <n> --output <out.json>
```

--starting-points was varied deliberately and is reported for every result; unless stated otherwise the batches used --starting-points 5. Every run in this study was a single-process invocation of the multiprocessor (MP) build. No run used HYPHYMPI, so the message-passing layer is excluded by construction rather than by argument.

The three options that materially change the fitted model are off by default and were therefore off in the main analysis: synonymous rate variation (--srv, default No), multinucleotide substitutions (--multiple-hits, default None), and the alignment-error absorbing rate class (--error-sink, default No). The first two correspond directly to the model-misspecification mechanisms of Wisotsky *et al.* (2020) and Venkat *et al.* (2018), and the third is directly relevant to alignments trimmed automatically, as ours are.

*Supports M2 (Reproducibility and convergence diagnostics)*

**Identical-command replication.** For each of a set of test genes, the identical command line (same alignment file, tree file, flags, binary and working directory) was executed repeatedly and the resulting $K$, log-likelihood and $P$ were compared across replicates (scripts/q2_replicates.py, scripts/seed_precheck.py). Replicates were run sequentially, not concurrently, so that no two replicates of the same gene competed for the same cores.

**Seed control.** HyPhy exposes a random seed through two syntaxes, an ENV="RANDOM_SEED=<n>" argument and a RANDOM_SEED environment variable. Both were tested at fixed values across replicates of the same command. In the extension the SetParameter batch-file form was tested under 2.5.100 and the ENV= form under 2.5.101 (Supplementary S2.12).

**Thread control.** The MP build parallelises with OpenMP and accepts CPU=<n>. To separate thread-scheduling non-determinism from other sources, three genes were run in a 2 × 4 design ({default, CPU=1} × four replicates) under otherwise identical commands (scripts/q2_replicates.py). The pre-registered extension repeated the CPU=1 condition, unseeded and seeded, on twenty genes (Supplementary S2.12).

**What was held constant, and to what precision.** For every paired comparison we verified that the input alignment and tree files were byte-identical, that the number of sequences and sites agreed, that the {Test} partition contained the same branches, and that the HyPhy binary and RELAX.bf version were the same. We also compared the nested fitting stages that precede the RELAX mixture model. The nucleotide GTR stage agrees between replicates to within $|\Delta$ log-likelihood$| \leq 2.7 \times 10^{-7}$, not exactly; we state the tolerance explicitly because "identical" would be false and because the residual is itself informative, being of the magnitude expected from a change in floating-point reduction order.

**Convergence flags and the re-fitting loop.** RELAX.bf re-fits an apparently failed comparison up to five times (while … loop_passes <= 5, L1529–1554); runs that exhaust those passes and still terminate with a negative likelihood-ratio statistic are recorded separately, because the alternative model nests the null and a converged fit therefore cannot be worse than it. A negative likelihood-ratio statistic is not a borderline result but an arithmetic impossibility under convergence, and requires no control, pairing or simulation to interpret.

**Why the intersection rule is stated.** An earlier internal version of one control script mixed denominators, using one arm's $n$ against the other arm's $n$, and produced a difference that was partly an artefact of unequal failure.

*Supports M2 (Null B)*

The pipeline was MAFFT protein alignment (Katoh and Standley, 2013), back-translation to codons, removal of columns with > 50 % gaps, and FastTree gene tree inference (Price *et al.*, 2010). 186574at4751 has a codon alignment that is not rectangular, one sequence being two nucleotides shorter than the rest, so tree inference fails and no run is possible; it was excluded on that ground alone, before any result was seen. No gene was selected or dropped on the basis of its result, so the set is the complete qualifying collection by construction. These are strongly constrained loci whose $\omega$ dynamics are not those of CAZymes; the property the null requires is not that they behave like CAZymes but that they have no reason to be relaxed *specifically on ECM branches*.

*Supports M2 (Null C, and the retired version of that control)*

Draws were made by scripts/gbe_batch1_refit.py, seed constant 20260730. The realised draws are deposited as labelled Newick files, one per gene, and it is those files, rather than the seed, that define the analysis: the drawing routine combined the seed constant with Python's built-in hash() of the gene name, which is salted per interpreter process, so re-running the script does not regenerate the same draws. We state this because it is a defect of exactly the kind this paper is about, and because the deposited trees make the analysis auditable regardless. Draws for the test-set-size ladder use a stable checksum instead and do reproduce from their recorded seed. All 88 draws were verified to contain zero ECM tips and a test set of size exactly 21. Alignments and trees were not recomputed, so the only difference from null B is which tips carry the label.

An earlier version of this control drew the 21 tips from all 51 genomes. Because 21 of the 51 are ECM, every draw then contained 5–13 ECM tips (mean 8.56, 41 %), and the control did not isolate the design contribution it was introduced to isolate. That version is retired; it is described here because its numbers appear in the preprint of this work.

*Supports M2 (codeml two-ratio comparison)*

The one-ratio model (model = 0) was compared against the two-ratio model (model = 2, the labelled branches as foreground) by a likelihood-ratio test on one degree of freedom; the two models are nested at the interior point $\omega_{fg} = \omega_{bg}$, so no boundary correction applies. Runs used NSsites = 0, F3x4 codon frequencies, an estimated transition/transversion ratio, cleandata = 0 so that the alignments are the ones RELAX analysed rather than a gap-stripped variant, and branch lengths estimated from scratch. Following our own recommendation about starting points, every model was fitted twice from $\omega = 0.5$ and $\omega = 1.5$ and the higher likelihood retained; the two starts also provide a paired reproducibility measurement in a second program, parallel to the starting-point comparison reported for RELAX.

Alignments were converted to sequential PHYLIP and the {Test} annotation to PAML's #1 foreground marker; the conversion was checked per gene by requiring the number of foreground labels to equal the number in the source tree and the number of tree tips to equal the number of sequences, and every gene passed. Each fit ran in its own working directory because codeml writes auxiliary files to the current directory and concurrent runs in a shared directory would overwrite one another silently. Of the 1,472 two-ratio fits across the four sets, none failed; the 1,152 quoted below for the truncation accounting is the first pass over the three original sets.

*Supports M2 (Clade Model C, and the truncation recovery pass)*

The alternative adds one parameter — a separate $\omega$ for the foreground branch set within one site class — and we confirmed on pilot genes that the fitted parameter counts differed by exactly one before interpreting any statistic on one degree of freedom.

These fits are far more expensive, and an initial pass did not complete. Each was capped at four hours and 161 of the 1,152 exceeded the cap, concentrated in the longest alignments and overwhelmingly at the higher starting value: 139 of the 576 fits begun at $\omega = 1.5$ did not finish, against 22 of the 576 begun at $\omega = 0.5$. Because the truncation rate differed between the observed and the simulated sets — 22 of 352, 69 of 400 and 70 of 400, or 6.3, 17.2 and 17.5 per cent — the sets would otherwise have rested on different denominators, and truncation is not random with respect to the outcome, since it discards the slowly converging fits. Every truncated fit was therefore re-run under a twenty-four-hour cap. Five of the observed-set fits were recovered in an earlier pass, leaving a final batch of 156, which completed in 30.4 hours of wall time with the longest single fit taking 14.0 hours; every one of the 161 finished, so the truncation was resource contention rather than non-convergence and the three sets are complete at 88, 100 and 100 genes. Where both starting values completed, the higher likelihood was retained, the same best-likelihood rule applied throughout. Every figure derived from these fits is reported with its denominator.

*Supports M2 (Clade Model C settings)*

Clade Model C was fitted as model = 3, NSsites = 2 with three site classes, against its null M2a_rel as model = 0, NSsites = 22 (Weadick and Chang, 2012). The two differ by exactly one parameter, so the likelihood-ratio test is on one degree of freedom. The divergence-matched set received its own recovery pass rather than the one described above for the other three sets.

*Supports M2 (Simulation positive control)*

Built and run by scripts/sim_positive_control.py and scripts/sim_relax_run.py; the divergence-matched rebuild by scripts/sim_divcal.py, scripts/sim_divcal_perK.py and scripts/sim_matched_gen.py.

**Generating parameters.** Each template supplied its three-class $\omega$ distribution and proportions from its RELAX null fit with $K$ fixed at 1, κ from its GTR fit, codon-site count from its alignment, and tree from its {Test}-labelled topology. Sequences were evolved under MG94 with F1x4 equilibrium frequencies. For the five-template set, 20 replicates were generated at $K = 1$ and 6 at each of $K = 0.5$ and 0.7. For the 78-gene panel the per-replicate seed is derived from gene, $K$ and replicate number, and realised pairwise distance was printed at generation as a self-check.

The bisection search was repeated separately at each generating $K$, since $\omega_{test} = \omega_{ref}^K$ accelerates the labelled branches whenever $\omega_{ref} < 1$ and a scale fitted at $K = 1$ therefore overshoots elsewhere. One template could not be brought to its target at any scale within the search bounds and was dropped. Only the branch-length scale is altered by this procedure; the $\omega$ classes, their proportions and κ are those of the first construction, so the generating value of $K$ is preserved. Realised identities are reported per replicate alongside their targets. Internal-branch labellings for the divergence-matched set reuse the same trees as the shallow internal arm, so the two differ in the alignments and not in the labelling (scripts/sim_relax_run2.py, which takes the tree path as an argument and reports the resolved source and label counts before launching).

The paired one- and five-starting-point fits serve two purposes: they supply the convergence check on data whose generating model is exactly the fitted model, and they measure the effect of the starting-point setting on type I error under a known null. This design answers three questions that the observational analysis cannot: whether the test rejects at its nominal rate when the null is exactly true ($K$ = 1); whether it recovers and detects genuine relaxation ($K$ = 0.5, 0.7); and whether simulated data pass the same convergence criteria that the real data fail — if they do, the criterion is falsifiable and the problem lies with the data, and if they do not, the problem lies with the optimiser.

*Supports M2 (Proportion conventions and aggregation)*

Two conventions for proportions run through the paper and are not the same family. Tests of a proportion against a fixed value are exact binomial; interval estimates of a proportion are Wilson score intervals. We name both because at the sample sizes involved the choice is not cosmetic: on the forty-family control of the companion study (Results 3) the Wilson interval is 22.1 to 50.5 per cent where the Clopper–Pearson interval is 20.6 to 51.7, and a reader recomputing from the counts alone would not recover our figures.

Aggregation of RELAX output used a tolerant JSON reader, since RELAX writes bare inf/-inf tokens that a strict parser rejects, silently dropping the gene.

### SUPPLEMENTARY S2 — SUPPORTING DETAIL FOR RESULTS 1 (THE FITTED VALUES ARE NOT REPRODUCIBLE, AND THE CAUSE IS THE OPTIMISER)

*S2.1 Nested fits in the eighty paired refits*

Supports the statement that the sub-stage fits preceding the RELAX mixture are recovered.

Nucleotide GTR log-likelihoods differed by more than 0.01 units in five of eighty pairs (median 0.0000; range -0.07 to +0.15), and MG94×REV log-likelihoods in eighteen of eighty (median 0.0000; range -0.38 to +1.99). One gene differed by one in its estimated parameter count.

*S2.2 Single-threaded execution: spread and cost*

Supports the statement that restricting HyPhy to one thread does not narrow the spread.

Under CPU=1 the log-likelihood range grew from 30.40 to 59.08 units in one gene and from 7.57 to 33.60 in another. Single-threaded execution is two to three times slower and buys nothing. The pre-registered extension (S2.12) repeated the single-threaded condition, two runs per gene, on twenty null orthologues. No gene returned the same log-likelihood twice: the within-gene ranges ran from 0.006 to 377 units (median 3.2), one gene fell within 0.01, and the range of $K$ within a gene ran from 0.0004 to 11.6. No $P$ value moved across 0.05 between the two runs. The nucleotide GTR log-likelihood agreed exactly as printed in fifteen of the twenty pairs, to within $8.4 \times 10^{-7}$ in four, and differed by $2.5 \times 10^{-3}$ in one; per-gene values are tabulated in S2.12.

*S2.3 The two fits issued with the same seed value*

Supports the statement that RANDOM_SEED has no effect on this analysis.

The two executions differ by 0.70 in $K$ and 63.5 in log-likelihood.

*S2.4 Design of the reduced-grid experiment*

Supports the statement that shrinking the Latin hypercube grid does not make a fit reproducible.

The program requires the starting grid to be at least ten times the number of starting points, which is why 10 is the smallest value it accepts. The design was three genes by three arms by four repeats, nine cells in all; one run failed, leaving eight cells with four repeats and one with three, and no cell produced two identical fits.

*S2.5 Repeat behaviour across HyPhy releases*

Supports the statement that the behaviour is not particular to the release we used.

Repeating the same command four times under HyPhy 2.5.28, 2.5.100 and 2.5.101 gave a distinct log-likelihood for every completed repeat in every gene under every version; one run under 2.5.28 ended in a segmentation fault, leaving that cell with three. The three releases span about five years and two generations of the analysis script, including the release in which the optimiser was rewritten with history-dependent heuristics and its line-search precision relaxed. We report the magnitudes as measured — median log-likelihood ranges of 48, 95 and 23 units in version order — but with three genes we do not read an ordering into them, and the comparison across the older version is one of repeatability within a version rather than of values between versions, since its analysis script differs.

*S2.6 The three kinds of failure, with per-gene values*

Supports the statement that the divergence between identical runs is not a single phenomenon.

Repeating an identical command produced divergent fits in every gene examined: two in the seed pilot, three in the threading control and twenty in the single-threaded extension (S2.12).

*Verdict reversal.* The seed example is the clearest case: the same command, twice, returns opposite conclusions about the same alignment.

*Catastrophic single-run failure.* In one gene, three of four identical executions settled within 28 log-likelihood units of one another (-9275.06, -9271.69, -9299.57), while the fourth landed 896 units worse (-10167.70). That worst fit returned $K = 1.000000$ exactly, with $P = 0.907$. A run that failed by the widest margin we observed reported the parameter value most easily read as "no departure from the null" — the boundary conceals the failure rather than revealing it.

*Unusable effect size under a stable verdict.* In a single-copy BUSCO gene with no expected signal, all eight repeated executions rejected the null (*P* between $7.5 \times 10^{-7}$ and $2.2 \times 10^{-3}$), all in the direction of intensification. Yet *K* ranged from 1.449 to 2.131, a spread of 47 per cent of the point estimate. Here the verdict is perfectly stable and the estimate is not; a study reporting only significance would record this gene as a reproducible finding.

*S2.7 PAML fits from two starting values of ω*

Supports the statement that the instability in PAML is confined to the models with site classes.

Within a four-hour limit, 22 of 576 fits initiated at $\omega = 0.5$ failed to complete, against 139 of 576 initiated at $\omega = 1.5$ (3.8 against 24.1 per cent), on alignments for which the alternative starting value converged within minutes. Terminations with a negative likelihood-ratio statistic were absent from all 368 two-ratio fits across the four sets and occurred in 2.3 per cent of site-class fits on the observed genes and 8.0 and 4.0 per cent on the two shallow simulated arms; on replicates matched to the divergence of the observed alignments the figure is 41.2 per cent.

*S2.8 Calibration at one against five starting points*

Supports the statement that more starting points make the calibration worse.

Among the ninety-four simulated replicates that converged under both settings, six moved from non-rejection to rejection and one moved the other way. The directional bias sharpened in step, from 87.5 to 95.2 per cent of rejections falling on the "relaxation" side, and the median estimate of *K* moved further from its true value of one.

*S2.9 GH15, fitted twice under an identical RELAX invocation*

Supports the statement that the transition from stable to unstable fits occurs at the mixture stage within a single analysis.

The nucleotide GTR stage agreed exactly; MG94×REV differed by $6 \times 10^{-4}$; the RELAX null and alternative models differed by 25.3 and 17.7 units in opposite directions, and the partitioned descriptive model by 40.2. The estimate of $K$ moved from 1.227 to 1.161 and the likelihood-ratio statistic from 114.6 to 28.7, a fourfold change with no change in the verdict.

*S2.10 The eighty paired refits at the RELAX stage*

Supports the statement that not one of the eighty null pairs reproduced its log-likelihood.

Of the eighty BUSCO null pairs fitted at one and at five optimiser starting points, thirty improved by more than 0.01 log-likelihood units at five and fifty worsened by more than 0.01; none agreed to within 0.01. The median change was -6.63 units and the range ran from -6,520 to +1,380. Because the Latin hypercube grid of initial values is drawn at random and was not held fixed, this is not a controlled test of --starting-points; what it establishes is weaker and sufficient, that the returned fit is not a function of the alignment and the user's settings alone. The corresponding comparison for the twenty-seven enzyme entries is Table 1.

*S2.11 Composition of the 410-fit convergence-flag census*

Supports the statement in Results 2 that the flag census is a census rather than a rate.

The pool comprises every RELAX fit in the observed analysis and its null distributions: 114 at one optimiser starting point on the observed alignments and the orthologue null, 88 for null A, 82 for null B, 81 for null C, 21 for the enzyme entries at five starting points, 13 for the ten-starting-point check, 6 for the entry re-fits and 5 for the starting-point ladder. The simulation, misspecification-control, test-fraction-ladder, CAZyme and replicate batches are analysed separately and are not in this pool.

Decomposed by what was fitted rather than by directory, the 410 are 350 null orthologues and 60 enzyme entries; by what was labelled, 175 carry a randomised test set and 235 the real ectomycorrhizal one. The directory holding the observed analysis is mixed, containing null B at one starting point alongside the entries, so a decomposition by directory alone would attribute 86 null-gene fits to the observed analysis, of which 33 are among the 184 that fired the flat-surface check. Of the 184 fits in which the flat-surface check fired, 162 are null orthologues and 22 enzyme entries; 87 carry a randomised labelling and 97 the real one. Regenerated by scripts/flag_provenance.py.

*S2.12 Why the documented seed has no effect, and what does*

Supports the statements in Results 1 that the seed does not reach the generator and that the analysis can nevertheless be made reproducible.

The command-line form ENV=RANDOM_SEED=<n> is parsed into a string held for later evaluation, and the startup routine that runs afterwards sets the seed from the clock and the process identifier and initialises the generator with it, then writes that value into the environment variable. The initialisation routine is called from exactly two places in the source tree: that startup path, and the branch of SetParameter that recognises an assignment to RANDOM_SEED. A command-line assignment reaches neither after startup, so the generator retains its clock-and-process seed while the variable reports the value the user supplied. We read this in the 2.5.100 sources and found the same two call sites, with the same ordering, in 2.5.28. The 2.5.101 sources add a third: after executing the stored command-line assignments, the startup routine now compares RANDOM_SEED against the seed in force and re-initialises the generator if they differ (global_things.cpp), so the command-line form seeds draws reproducibly there — verified by the two-run test below, which returns identical variates under 2.5.101 and different ones under 2.5.100. The release notes of 2.5.101 do not mention the change. Every analysis in this manuscript ran under 2.5.100.

The behaviour reproduces in a few seconds without any data. Under 2.5.100, a batch file drawing five variates and printing RANDOM_SEED alongside them, run twice as hyphy ENV=RANDOM_SEED=12345, reports the seed as 12345 both times and returns different variates; the same file with SetParameter (RANDOM_SEED, 12345, 0) prepended returns identical variates on every run.

Applied to RELAX, the two available measures were tested against each other on three of the null orthologues. Seeding through SetParameter while holding the thread count at one returned a single distinct log-likelihood in each of six gene-by-setting combinations, twenty-one runs in total: three genes repeated four times at one optimiser starting point (-9297.2827396846, -13117.9963255417 and -13206.0795973072) and the same three repeated three times at five (-9287.8855692030, -13116.9679781510 and -13196.2645335656), with $K$ likewise constant and the output files differing only in their timing fields. Neither measure worked alone. Holding the thread count at one without seeding left all seven repeat pairs divergent, by 2.36 to 296.61 log-likelihood units, and moved the $P$ value across 0.05 in three of them: from 0.417 to 0.0032, from 0.0629 to 0.0125, and from $8.0 \times 10^{-8}$ to 0.78. Seeding without holding the thread count narrowed one gene from 7.84 log-likelihood units to 0.048 but did not close it.

A pre-registered extension, with gene list, arms, criterion and predictions fixed before any run, took the comparison to twenty null orthologues: the three above and seventeen drawn at random, with a recorded seed, from the remaining eighty-six, single-copy BUSCO genes of 50 or 51 sequences, every fit at one optimiser starting point. Four arms were run. Under 2.5.100, the stock binary at one thread, two runs per gene, and the SetParameter seed at one thread, three runs per gene; under 2.5.101, the documented command-line seed at one thread, three runs per gene, and the same seed at the default thread count, three runs each of five genes. The criterion was that of the three-gene runs, every run of a gene returning the same alternative-model log-likelihood as printed. The two seeded single-thread arms reproduced all twenty genes, sixty runs each, *K* and *P* likewise identical within gene. The stock single-thread arm reproduced none: the within-gene log-likelihood ranges ran from 0.006 to 377 units with a median of 3.2, one gene fell within 0.01, and no *P* value moved across 0.05. The seeded default-thread arm reproduced none of its five genes, one gene returning the same fit in two of its three runs, with ranges of 0.29 to 107 units and one *P* value moved across 0.05. All four outcomes were as predicted. In the two arms that did not reproduce, the boundary value $K = 1.0000$ exactly, with $P = 1$, appeared in three of forty and two of fifteen runs, and in every case that run carried the worse log-likelihood of its gene, by 2.9 to 107 units. The nucleotide GTR stage agreed exactly as printed in fifteen of the twenty single-thread pairs, to within $8.4 \times 10^{-7}$ in four, and differed by $2.5 \times 10^{-3}$ in one. Per-gene ranges follow; a dash marks a gene not in the five-gene arm.

| **Gene** | **Stock 2.5.100, one thread, two runs: Δ log-likelihood** | **Δ *K*** | **Δ GTR log-likelihood** | **Seeded 2.5.101, default threads, three runs: Δ log-likelihood** | **Δ *K*** | **Distinct log-likelihoods** |
|---|---|---|---|---|---|---|
| 123228 | 36.9 | 0.00501 | 0.00247 | — | — | — |
| 148293 | 67.8 | 0.0876 | 0 | — | — | — |
| 167374 | 24.1 | 0.217 | $8.4 \times 10^{-7}$ | — | — | — |
| 176648 | 174 | 0.0393 | 0 | 0.456 | 0.0206 | 2 of 3 |
| 176748 | 2.96 | 0.0559 | 0 | — | — | — |
| 192135 | 0.376 | $3.9 \times 10^{-4}$ | 0 | — | — | — |
| 202089 | 319 | 11.6 | 0 | — | — | — |
| 20952 | 0.373 | 0.0081 | 0 | — | — | — |
| 245480 | 12.8 | 0.0212 | 0 | 0.291 | 0.00518 | 3 of 3 |
| 316920 | 0.0548 | 0.0213 | 0 | — | — | — |
| 322267 | 2.97 | 0.00536 | 0 | — | — | — |
| 332416 | 2.34 | 0.0208 | 0 | — | — | — |
| 398592 | 0.00591 | 0.00273 | 0 | — | — | — |
| 428342 | 2.9 | 0.0568 | 0 | 67.8 | 0.612 | 3 of 3 (*P* |

| | | | | | | |
|---|---|---|---|---|---|---|
| | | | | | | crosses 0.05) |
| 428984 | 0.215 | 0.00629 | 0 | — | — | — |
| 438731 | 21.8 | 0.103 | 0 | — | — | — |
| 452573 | 0.564 | 0.201 | 0 | 0.489 | 0.0298 | 3 of 3 |
| 453693 | 6.5 | 0.0211 | $1.1 \times 10^{-7}$ | 107 | 0.0499 | 3 of 3 |
| 54252 | 377 | 0.0896 | $1.7 \times 10^{-7}$ | — | — | — |
| 79797 | 3.35 | 0.0208 | $3.2 \times 10^{-7}$ | — | — | — |

The three-gene runs used HyPhy 2.5.100 on single-copy orthologues of 51 sequences, and the extension 2.5.100 and 2.5.101 on twenty of 50 or 51; we did not test 2.5.28 in either, nor deeper alignments, multi-copy families, or more than one starting point in the extension. Regenerated by rng_build/{pilot_run.sh, cpu1_run.sh, main_run.sh, sp5_run.sh} and read by scripts/seedfix_verdict.py into logs/RESULT_FINAL_seedfix.txt, and for the extension by scripts/repro20_launch.py and scripts/repro20_verdict.py into logs/RESULT_FINAL_repro20.txt; the few-second demonstration is rng_build/mre/run_mre.sh.

*S2.13 Reporting practice in studies that use RELAX*

Whether the failures documented above are visible to a reader of the literature is an empirical question, and we asked it of the papers that use the program. Europe PMC was searched over full text for ("RELAX" AND "HyPhy" AND "selection") on 28 August 2026. Because RELAX is also an ordinary English word, a record was counted as using the program only where the token appeared with a word boundary alongside at least one of HyPhy, Wertheim, selection intensity, relaxed selection or intensified selection in the same document.

The search returned 321 records, of which 305 carried a PubMed Central identifier and were requested; 268 full texts were retrieved and 37 returned HTTP 404, and 34 of those retrieved failed the co-occurrence filter, leaving a denominator of 234 papers examined. The 37 records whose full text could not be retrieved were not examined and are not represented below. The index this query runs against grows, so the figures are a snapshot rather than a fixed set; the same query a week earlier, on 21 August 2026, returned 319 records and a denominator of 230, and gave the same zero.

Of those 234, none reported a random-number seed. Two (0.9 per cent) reported the number of optimiser starting points, and between three and five (1.3 to 2.1 per cent) described repeating a fit in order to check that it reproduced rather than for some other purpose. The regular expressions match more broadly than that — eleven papers and twenty-three respectively — and the difference is read out by hand: most of the starting-point matches are the ordinary English phrase, and most of the repetition matches are bench replicates or statements that a trait evolved several times. The two figures we report are the matches that survive reading the surrounding sentence, and the range on the second is the three that state the purpose explicitly against the two further papers that repeat a fit without saying why.

Three checks stand behind the zero, and it moved twice before settling. An earlier pass over a narrower denominator returned two matches for a seed; read in context both were false positives, one a random seed of samples in an expression analysis and the other a seed supplied to a different program. Widening the filter to recover papers that first pass had missed, and removing those two patterns, returned no matches at all. A positive control runs on every execution: the present manuscript, which states all three of the items the patterns look for, is passed through the same patterns and must match, so a pattern that had silently stopped working would halt the script rather than return a zero.

Two further counts produced by the same script are deliberately not reported. A figure for how often the HyPhy version is stated is inflated by the program's citation title appearing in reference lists, and a figure for code or data deposition is dominated by links to other authors' repositories;

both would need redefining before they could be read. A count of papers mentioning convergence does not separate a diagnostic from a passing use of the word.

Regenerated by c_audit2.py, which writes RESULT_FINAL_caudit.txt: the accounting above, the identifiers of all 234 papers in the denominator, and the matched sentence for every hit, so that the reading behind each reported figure can be repeated.

### SUPPLEMENTARY S3 — SUPPORTING DETAIL FOR DISCUSSION 2 (WHAT TO DO INSTEAD)

*S3.1 Starting-point ladders beyond the two-configuration case*

Supports the recommendation to fit from at least three configurations and report the spread.

In a second family the likelihood improved monotonically from one to five to ten starting points and then collapsed by 1,873 units at twenty. In a third, the best-fitting run of four was the one that returned no significant result. Whichever configuration a study happens to run is the configuration whose answer gets published.

### SUPPLEMENTARY TABLE S4

**The 51 genomes carrying the codon alignments used in every selection analysis.** These are the species with coding sequences recoverable in a form suitable for codon-aware alignment; they are a subset of the 183-genome annotation panel and of the 182-tip phylogenetic panel, and they are not a random subset of either (Discussion 3). "Lifestyle (used here)" is the assignment under which every selection analysis and every null set in this paper was built. "Lifestyle (corrected)" is filled only for the three genomes the annotation audit of the companion study (Results 2) reassigned, and shows what they would be under the corrected assignment; the selection analyses were not re-run on it, and Methods M2 states why. Under the assignment used here the test set is 21 ectomycorrhizal tips of 51; under the corrected one it would be 18. "Public" records whether the genome was publicly released at the time of analysis: 35 of the 51 were, and 16 were not. Portal codes are the stable JGI MycoCosm identifiers and are sufficient to retrieve each genome.

| Portal | Species | Lifestyle (used here) | Lifestyle (corrected) | Public |
|---|---|---|---|---|
| Altal1 | Alternaria alternata SRC1lrK2f v1.0 | Pathogen | — | yes |
| Amagr1 | Amanita aff. grandis KIS10 v1.0 | Ectomycorrhizal | — | **no** |
| Amamu1 | Amanita muscaria Koide v1.0 | Ectomycorrhizal | — | yes |
| Amapyr1 | Amanita aff. conicoverrucosa KIS12 v1.0 | Ectomycorrhizal | — | **no** |
| Amath1 | Amanita thiersii Skay4041 v1.0 | Saprotroph | — | yes |
| Amore1 | Amorphotheca resinae v1.0 | Saprotroph | — | yes |
| Ascim1 | Ascobolus immersus RN42 v1.0 | Saprotroph | — | yes |
| Aurde3_1 | Auricularia subglabra v2.0 | Wood decayer | — | yes |
| Aurpu_var_pul1 | Aureobasidium pullulans var. pullulans EXF-150 | Saprotroph | — | yes |
| Bissp1 | Bisporella sp. PMI_857 v1.0 | Saprotroph | — | yes |
| Bolcoc1 | Boletus coccyginus 2016PMI039 v1.0 | Ectomycorrhizal | — | **no** |
| Botbo1 | Botryobasidium botryosum v1.0 | Saprotroph | — | yes |
| Cananz1 | Cantharellus anzutake C23 v1.0 | Ectomycorrhizal | — | yes |

| | | | | |
|---|---|---|---|---|
| Cenge3 | Cenococcum geophilum 1.58 v2.0 | Ectomycorrhizal | — | yes |
| Chove1 | Choiromyces venosus 120613-1 v1.0 | Ectomycorrhizal | — | yes |
| ClaPMI390 | Clavulina sp. PMI_390 v1.0 | Saprotroph | — | yes |
| Conpu1 | Coniophora puteana v1.0 | Wood decayer | — | yes |
| Dacsp1 | Dacryopinax primogenitus DJM 731 SSP1 v1.0 | Wood decayer | — | yes |
| ElagrMar1 | Elaphomyces granulatus Maridalen v1.0 | Ectomycorrhizal | — | **no** |
| Fibsp1 | Fibulorhizoctonia sp. CBS 109695 v1.0 | Saprotroph | — | yes |
| Gloin1 | Rhizophagus irregularis DAOM 181602 v1.0 | Arbuscular mycorrhizal | — | yes |
| Gyman1 | Gymnopus androsaceus JB14 v1.0 | Saprotroph | — | yes |
| Hebcy2 | Hebeloma cylindrosporum h7 v2.0 | Ectomycorrhizal | — | yes |
| Hetan2 | Heterobasidion annosum v2.0 | Pathogen | — | yes |
| Hyafin1 | Hyaloscypha finlandica PMI_746 v1.0 | Ericoid mycorrhizal | — | **no** |
| Hyavar1 | Hyaloscypha variabilis MD1 v1.0 | Ericoid mycorrhizal | — | **no** |
| Hypsu1 | Hypholoma sublateritium v1.0 | Wood decayer | — | yes |
| Hyssto1 | Hysterangium stoloniferum HS.BST v1.0 | Ectomycorrhizal | — | yes |
| Jaaar1 | Jaapia argillacea v1.0 | Wood decayer | — | yes |

| | | | | |
|---|---|---|---|---|
| Kurarg1 | Kurtia argillacea OMC1749 v1.0 | Ectomycorrhizal | Ericoid mycorrhizal | **no** |
| Lacaka1 | Lactarius akahatsu QP v1.0 | Ectomycorrhizal | — | yes |
| Lacbi2 | Laccaria bicolor v2.0 | Ectomycorrhizal | — | yes |
| Matter1 | Mattirolomyces terfezioides MAT_tt4AIII v1.0 | Ectomycorrhizal | — | **no** |
| MelPMI1271_1 | Meliniomyces sp. PMI_1271 v1.0 | Ericoid mycorrhizal | — | **no** |
| Melbi2 | Meliniomyces bicolor E v2.0 | Ericoid mycorrhizal | — | yes |
| Morco1 | Morchella importuna CCBAS932 v1.0 | Ectomycorrhizal | Saprotroph | yes |
| Neole1 | Neolentinus lepideus v1.0 | Wood decayer | — | yes |
| Neucr_trp3_1 | Neurospora crassa FGSC 73 trp-3 v1.0 | Saprotroph | — | yes |
| Paxin1 | Paxillus involutus ATCC 200175 v1.0 | Ectomycorrhizal | — | yes |
| Penswi1 | Penicillium swiecickii 182_6C1 v1.0 | Ectomycorrhizal | Saprotroph | **no** |
| Phchr2 | Phanerochaete chrysosporium RP-78 v2.2 | Wood decayer | — | yes |
| Phlbr1 | Phlebia brevispora HHB-7030 SS6 v1.0 | Wood decayer | — | yes |
| Ruscya1 | Russula cyanoxantha BavarianF04 v1.0 | Ectomycorrhizal | — | **no** |
| Ruseme1 | Russula emetica Přilba v1.0 | Ectomycorrhizal | — | **no** |
| Rusoch1 | Russula ochroleuca Přilba v1.0 | Ectomycorrhizal | — | **no** |
| Sebve1 | Sebacina vermifera MAFF 305830 v1.0 | Orchid mycorrhizal | — | yes |

| | | | | |
|---|---|---|---|---|
| Ser400_1 | Serendipita sp. 400 v1.0 | Orchid mycorrhizal | — | **no** |
| Ser405_1 | Serendipita sp. 405 v1.0 | Orchid mycorrhizal | — | **no** |
| Serend1 | Serendipita sp. 407 v1.0 | Orchid mycorrhizal | — | **no** |
| Thacu1 | Rhizoctonia solani AG-1 IB | Pathogen | — | yes |
| Theter1 | Thelephora terrestris UH-Tt-Lm1 v1.0 | Ectomycorrhizal | — | yes |

Penswi1 was recorded at the time of download as *Penicillium* sp. GLBRC 242 v1.0, which is a different assembly (31.35 Mb, 11,517 gene models); the files analysed here are the 34.13 Mb, 12,802-model assembly of *Penicillium swiecickii* 182_6C1 v1.0, and the name in this table is corrected accordingly. Its pre-audit lifestyle label, ectomycorrhizal, was inherited from that other assembly's portal record rather than read from the record of Penswi1 itself, so it is not evidence that the portal assigns *P. swiecickii* to ectomycorrhizal fungi. The lifestyle audit and every count are unaffected, since they were made on the files.

The count analyses do not depend on the unpublished genomes: restricting the primary contrast to public genomes leaves the estimate essentially unchanged (companion study, Results 2). The selection analyses do depend on them, and cannot be reproduced from public data alone.
